# Post-Decision Gaze as a Behavioral Manifestation of Decision Confidence

**DOI:** 10.1101/2024.08.28.610145

**Authors:** Yonatan Stern, Ophir Netzer, Morris Goldsmith, Danny Koren, Yair Zvilichovsky, Uri Hertz, Roy Salomon

## Abstract

Decisions are naturally accompanied by a feeling of confidence in their correctness. Typically measured via self-reported judgments, decision confidence is also expressed implicitly through sensorimotor behavior. In one exploratory and two preregistered experiments (total *N* = 115), we combined VR-based probabilistic-learning and spatial-cueing paradigms with computational modeling to examine whether post-decision gaze can serve as an implicit behavioral measure of decision confidence. Gaze direction generally aligned with predictive decisions about a target’s upcoming location, while the degree of alignment tracked the model-based probability underlying the decision. Gaze–decision alignment also exhibited previously established statistical hallmarks of confidence. Under increased uncertainty, gaze and decisions tended to diverge, reflecting strategic “hedging” behavior. Gaze–decision alignment and explicit confidence judgments were modestly correlated, and remained correlated after controlling for the decision’s probability, suggesting that the two are partially overlapping manifestations of internal confidence. Notably, reported confidence showed higher metacognitive sensitivity and increased serial dependence across trials, while gaze-based confidence expressed a more trial-specific readout of internal confidence. These findings suggest that post-decision gaze in visual-spatial learning tasks is an implicit, embodied manifestation of confidence. The complementary functional characteristics of implicit and explicit expressions of confidence may support adaptive behavior during learning.

## 1. Introduction

Our decisions are usually accompanied by an internal sense of confidence in their correctness. Confidence is the product of second-order, metacognitive processes that evaluate first-order performance on cognitive tasks and decisions. Importantly, several theoretical accounts propose that confidence is computed by integrating multiple informational cues, including post-decisional, motor, and physiological signals, rather than reflecting a simple readout of evidence quality (e.g., Fleming & Daw, 2017; Kiani & Shadlen, 2009; Koriat, 2012). Thus, confidence in a decision may reflect different types and combinations of cues, depending on when and how it is computed.

The standard experimental method for directly probing this internal sense of confidence is to elicit an explicit judgment or rating of one’s level of confidence (e.g., Maniscalco & Lau, 2012; Rahnev et al., 2020). However, the degree of overlap between such explicit judgments and the actual level of internal confidence has been increasingly questioned. First, confidence judgments are influenced by interoceptive and sensorimotor processes and information that extend beyond the decision itself (Allen et al., 2016; Kiani et al., 2014). Whether internal confidence (e.g., the signal that might drive behavioral expressions) is similarly influenced by such factors is currently unknown. Second, prompted confidence judgments may also introduce and be affected by additional evaluation and response processes that do not occur otherwise (Shekhar & Rahnev, 2021). In contrast, various sensorimotor processes that occur naturally and spontaneously during the first-order decision process have also been shown to reflect a second-order evaluation of that process in an implicit and unintrusive manner. Most prominently, reaction time is an often-used implicit index of internal confidence, exhibiting robust, moderately sized trial-level correlations with explicit confidence ratings (Rahnev et al., 2020; Weidemann & Kahana, 2016). More recently, additional indices such as motor adaptation (Dotan et al., 2018; Faivre et al., 2020), speech prosody (Goupil et al., 2021), and pupil dilation (Colizoli et al., 2018; Urai et al., 2017) have been shown to reflect internal confidence in ways that are tightly coupled to ongoing behavioral regulation. Thus, implicit sensorimotor manifestations of internal confidence also help reveal the role of confidence in regulating action and social interaction, capturing additional, functional aspects of confidence that may remain untapped by explicit metacognitive judgments (Faivre et al., 2020; Goupil et al., 2021).

Confidence is critical for adaptive functioning in a dynamic environment (Fleming, 2024; Nelson, 1984), guiding a wide range of other types of everyday behavior (Goldsmith & Koriat, 2007), such as learning and information-seeking behavior (Desender et al., 2018; Drugowitsch et al., 2019). Although historically, the study of confidence and metacognition has focused primarily on the domains of perceptual decisions and memory (Fleming, 2024; Rahnev et al., 2020), interest in decision confidence during learning has grown in the past decade. During probabilistic learning, confidence judgments track a decision’s underlying likelihood distribution (Meyniel et al., 2015) and reflect the crossing of an internal decision boundary when choosing between competing options (Hertz et al., 2018).

In line with the adaptive role of confidence in representing and signaling the assessed accuracy of one’s decisions, confidence ratings robustly distinguish correct and incorrect decisions, and humans show significant metacognitive sensitivity (Maniscalco & Lau, 2012; Rahnev et al., 2020). However, confidence judgments also diverge from the first-order decision process in significant ways. They integrate priors more optimally (Constant et al., 2023) and include parallel or additional post-decision computational processes (Balsdon et al., 2020; Fleming & Daw, 2017). Confidence judgments also exhibit biases such as “leakiness,” with judgments from previous trials continuing to exert an influence on later judgments, even across tasks and domains (Mei et al., 2023; Rahnev et al., 2015). Research focusing on confidence reports has extensively characterized confidence judgments; however, it remains an open question whether sensorimotor behavior that reflects a second-order assessment of a decision exhibits similar characteristics.

Despite extensive research on confidence and sensorimotor behavior, and the close connection between gaze behavior and cognitive processes underlying decisions (Spering, 2022), the relation between gaze and confidence in humans remains surprisingly underexplored. Gaze is typically directed toward features and regions of the environment perceived as important (Krajbich et al., 2010; Niv, 2019; Spering, 2022). For example, in viewing naturalistic scenes, the learned statistical regularities of scene composition guide attention and visual search, enabling efficient scanning, filtering and processing of complex visual input (for review, see Wolfe, 2020). Thus, gaze is shaped by one’s experience, providing a window to an internal model of the environment that one has learned (Bar, 2004; Leong et al., 2017). Examining gaze’s relation to the evaluation of decisions, non-human primate studies have demonstrated that gaze metrics reflect both decision certainty (Seideman et al., 2018) and the reevaluation of one’s decision during learning (Ferro et al., 2024). Although decisions and ocular behavior often occur in tandem, previous work in humans has largely examined them separately, leaving the nature of their interaction and contribution to adaptive behavior unclear.

The current study examines whether post-decision gaze can serve as an implicit, behavioral manifestation of decision confidence during probabilistic learning. Throughout the manuscript, we distinguish between three related but non-identical constructs. First, decision certainty refers to the model-based probability underlying the decision made during the learning task. Second, internal confidence refers to the second-order evaluation of the decision, which may depend on, but is not reducible to decision certainty (Pouget et al., 2016). Third, confidence expressions refer to the behavioral manifestations of internal confidence, including explicit confidence reports and the present measure derived from gaze behavior. The specific metric of post-decision gaze we examine is the degree of alignment between a participant’s gaze and their decision’s direction. This gaze-based measure is inspired by post-decision wagers that are used to assess metacognitive processes in humans (Koren et al., 2006; Persaud et al., 2007), as well as measures such as ‘waiting time’ and ‘looking time’ that serve as behavioral proxies for confidence in animal studies (Kepecs & Mainen, 2012; Schmack et al., 2021). Like explicit confidence judgments, these measures are second-order in the sense that they are either explicitly (wagering) or implicitly (waiting and looking time) contingent upon the decision (Fleming et al., 2012), and are modulated by decision certainty (Kepecs & Mainen, 2012).

In three experiments, participants performed a probabilistic learning task in which they predicted the location (left or right) of an upcoming visual target based on probabilistic cues, and immediately afterwards reached for the target when it appeared inside a virtual-reality environment. These left-right predictions are also referred to as *decisions*. Computationally modeling the latent probability underlying the decision made on each trial (i.e., *p*(Choice *_t_*)) served as an index of decision certainty during learning. We also measured trial-by-trial fluctuations in the degree to which post-decision gaze was directed toward or away from the predicted target. Focusing on changes in gaze–decision alignment, we examined their association to decision certainty indexed by the model-based decision probability, as well as to the explicit confidence ratings elicited in Experiment 3. Specifically, we asked: (1) Does the degree of alignment between post-decision gaze and visual-spatial decisions implicitly express a person’s internal confidence in those decisions? (2) If so, how does gaze—decision alignment relate to reported confidence judgments and how do their functional characteristics compare? (3) How do fluctuations in gaze—decision alignment support adaptive behavior in a dynamic environment?

In two initial experiments (one exploratory, one preregistered), the overall binary direction (left or right) of post-decision gaze generally converged with the predicted target location. More importantly, a continuous measure of gaze–decision alignment was found to track trial-by-trial variation in the model-based decision probability that indexes decision certainty, while also exhibiting the three previously established statistical hallmarks of confidence (Sanders et al., 2016). In a third experiment, gaze–decision alignment correlated modestly at the trial level with self-reported confidence ratings, even after controlling for decision probability. Together, these findings suggest that in the present visual-spatial paradigm, gaze–decision alignment constitutes an implicit, behavioral manifestation of decision confidence that partially overlaps with explicitly reported confidence. A manifestation we refer to as ‘gaze-based confidence.’ Comparing gaze-based and reported confidence’s functional characteristics, reported confidence exhibited superior metacognitive sensitivity, while gaze-based confidence was less influenced by the level of confidence on preceding trials, reflecting a more momentary readout. Illustrating the potential functional benefits of two partially overlapping, complementary measures, whereas reported confidence decreased with decreased decision certainty, the direction of post-decision gaze increasingly diverged from the predicted target direction, serving to “hedge” one’s decision under uncertainty.

## 2. Results

### 2.1 Participants learn the probabilistic rule

We first report the results of Experiments 1 and 2, followed by Experiment 3, which also included explicit confidence ratings. We confirmed that participants’ decisions exhibited the well-established signatures of probabilistic learning. Per our preregistration, we define accuracy as whether predictions conformed to the underlying mapping rule (“rule accuracy”). This is distinct from predicting the target’s actual location, which is constrained by the 75% cue-validity. Across both experiments, rule accuracy was significantly higher than chance (*M* _accuracy_ = 71.5%, 95% CI = [69.7, 73.3], *t*_110_ = 23.5, *p* < .001, Cohen’s *d* = 2.2; see Fig. 1B, Fig. S1, and SM for similar results when analyzing actual accuracy and individual experiments). In addition, accuracy within blocks of trials governed by a single rule exhibited a classic learning-curve shape (see Fig. 1C and Fig. S2). Thus, participants were engaged and able to learn the underlying probabilistic mapping rule.

**Figure 1.**
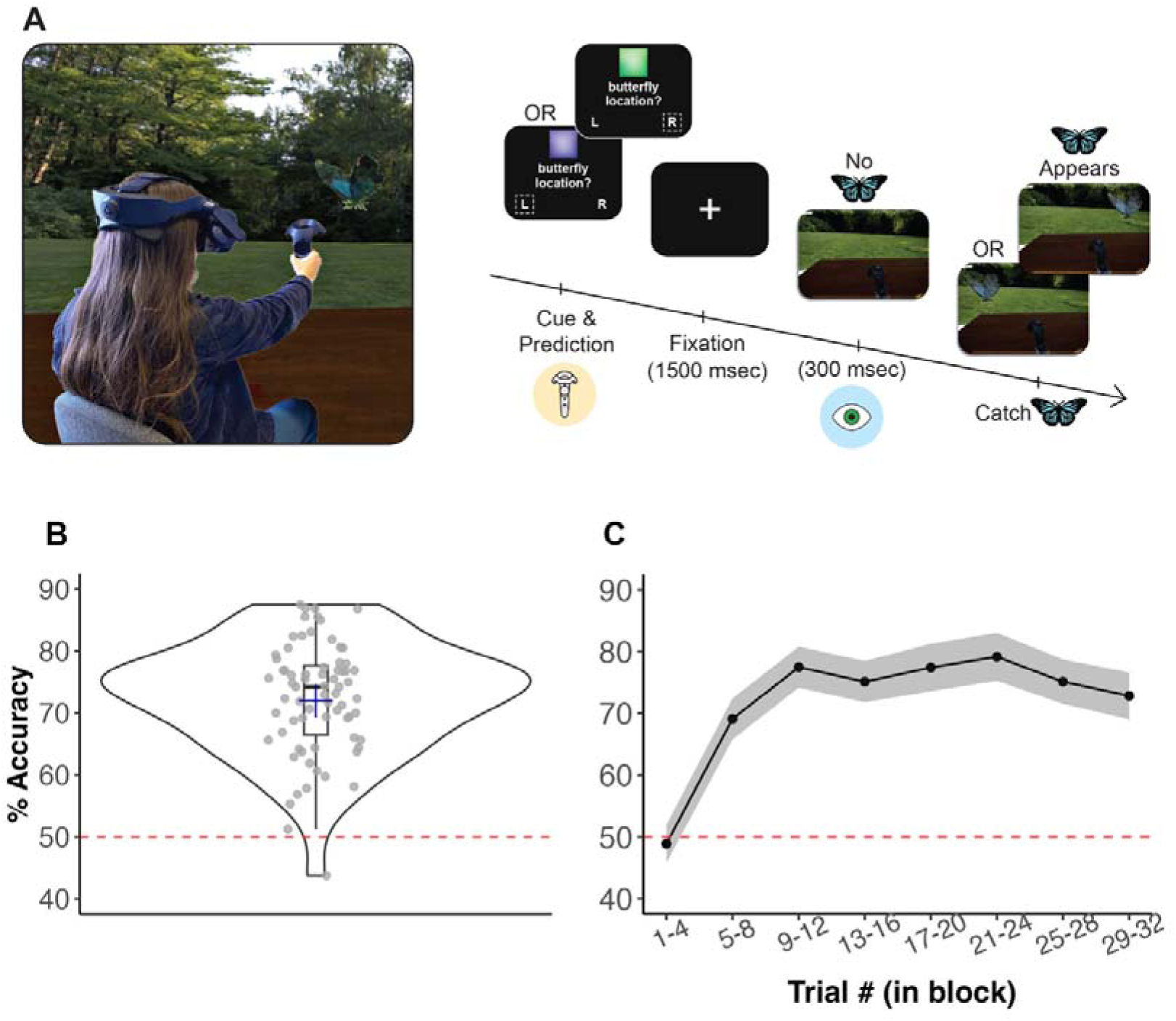
Experimental paradigm. **Panel A:** (Left) Experimental setup. Participants sat at a table and reached toward the target butterfly when it appeared inside the virtual-reality (VR) environment. (Right) Trial Flow. A colored cue prompted participants to predict the upcoming target’s location (Right/Left) via the controller’s touchpad, with no display change post-response. Following the response, a fixation cross preceded entry into the virtual environment, where, after 300 milliseconds, a target appeared. Horizontal gaze direction was measured during this 300 milliseconds interval. After which a butterfly appeared in one of six static locations on the right or left side of the environment. Following the butterfly’s appearance, participants were instructed to reach toward it as quickly and accurately as possible. **Panels B–C: Participants’ decisions exhibited classical probabilistic learning signatures across Experiments 1 and 2.** Given our interest in participants’ learning of the underlying cue-location mapping rule, we define accuracy as whether the response conformed to the underlying probabilistic rule. **(B) Accuracy exceeded chance**. Gray dots denote individual participants (including those subsequently excluded by preregistered criteria), the dashed red line indicates chance level (50%), and the blue cross represents the group mean. **(C) Learning curve within blocks.** Averaging trials across blocks, accuracy was low at the start of the block, and quickly increased as participants learn the underlying rule until a plateau was reached. Black line is group average and gray area represents the 95% confidence interval across participants.

### 2.2 Gaze converges with predictions and tracks trial-by-trial decision certainty

Gaze Direction, calculated as the time-weighted average of the normalized horizontal gaze eccentricity during the 300 millisecond interval preceding target onset, differed significantly between trials with Right versus Left predictions, reflecting an overall correspondence between predictions and subsequent gaze direction (*Exp. 1*: *M* _normalized gaze difference_ = 0.95, 95% CI= [0.80, 1.10], *t*_24_ = 660.77, *p* < .001, Cohen’s *d* =2.56; *Exp. 2*: *M* _normalized gaze difference_ = 0.90, 95% CI= [0.79, 1.01], *t*_24_ = 922.83, *p* < .001, Cohen’s *d* =3.04; see Fig. 2A). This difference was highly consistent and statistically significant for 54 of 56 participants at the individual level (see Fig. S3). Likewise, the binarized Gaze Direction measure (i.e., Right/Left, based on the sign of Gaze Direction) converged with the predicted direction in 73% of trials (95% CI = [70%, 75%]; see Fig. 2B and Fig. S4). Thus, as expected, the overall direction of post-decision gaze was largely aligned with the participants’ preceding, explicit prediction.

**Figure 2.**
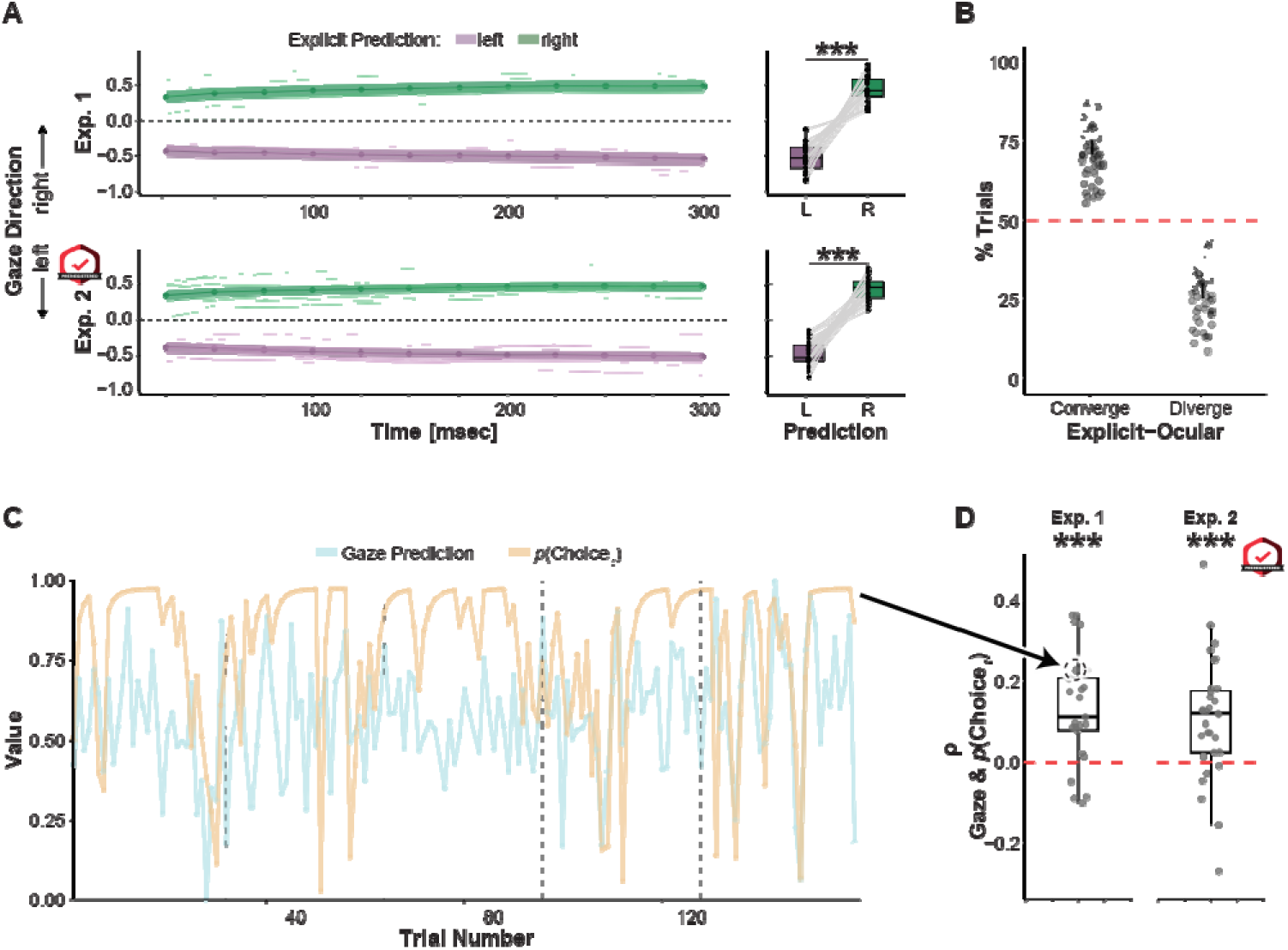
Convergence of predictions and post-decision gaze. **Panel A: Gaze Direction is aligned with the prediction’s direction.** (Left) Time course of normalized time-weighted post-decision gaze direction in the 300 milliseconds interval prior to target appearance, averaged across participants. (Right) Mean normalized gaze direction averaged across participants and trials is robustly aligned with the predicted location. **Panel B: Convergence and divergence of explicit prediction’s and gaze’s direction across participants**. Gaze was binarized into a Right/Left direction analogous to explicit predictions. Across both experiments, gaze’s direction and the prediction’s direction converged in ∼70% of the trials. The black line reflects a 95% confidence interval, and gray dots represent individual participants. **Panel C: Correlation between trial-by-trial Gaze Prediction and model-based probability of the decision *p*(Choice *_t_*).** A random participant’s trajectory across the experiment of *p*(Choice *_t_*), the probability of making the decision derived from the computational model, and Gaze Prediction, gaze magnitude normalized by the prediction’s direction. Note that for the clarity of the visualization, Gaze Prediction values were bound to the limits of *p*(Choice *_t_*) between 0 and 1. **Panel D: Distribution of individual participants’ correlation coefficients between *p*(Choice *_t_*) and Gaze Prediction.** In both experiments the coefficients’ distributions were significantly positive. Thus, post-decision gaze direction was more tightly aligned with the prediction’s direction as the underlying probability of the prediction increased.

We next examined whether trial-by-trial fluctuations in the degree of alignment between gaze direction and explicit predictions track variation in the decision certainty as measured by the model-based probability of the decision made (i.e., (*p*(Choice *_t_*)). We utilize Gaze Prediction, that is Gaze Direction adjusted by the explicit decision’s direction, providing a continuous measure of gaze–prediction alignment (see Methods and Materials). While we initially preregistered the use of a simple Rescorla-Wagner (RW) learning model, formal model comparison (see Fig. S5) led us to choose a better-fitting RW model that includes a stickiness parameter (Culbreth et al., 2016; Daw et al., 2011; see SM for similar results with the simple RW model). Following our preregistration analysis plan, the trial-by-trial correlation between *p*(Choice *_t_*) and Gaze Prediction was calculated per participant (see Fig. 2C). The group distribution of Pearson correlation coefficients was significantly greater than zero in both experiments (*Exp. 1*: Mean = .13, 95% CI = [.08, .19], *t*_24_ = 4.84, *p* < .001, Cohen’s *d* = 0.97, *Exp. 2*: Mean = .11, 95% CI = [.05, .16], *t*_24_ = 4.02, *p* = .001, Cohen’s *d* =0.82; Fig. 2D and Fig. S6). Thus, the greater the decision certainty as indexed by the model-based probability underlying the decision on a given trial, the more tightly post-decision gaze aligned with the predicted target direction.

We further examined whether the relationship between gaze and decision certainty indexed by *p*(Choice *_t_*) is graded or driven solely by “divergent” trials in which the gaze and decision directions differed. We reexamined the correlation between Gaze Prediction and the model-based decision probability while controlling for the binary component of alignment (coded as -1 or +1 for diverging and converging trials). Across both experiments, the combined group-level partial correlation remained significantly greater than zero (Mean = .07, 95% CI = [.04, .10], *t*_55_ = 4.94, *p* < .001, Cohen’s *d* = 0.66). Thus, after controlling for the binary component of the gaze—decision relationship, trial-by-trial fluctuations in the graded alignment between gaze and predicted direction tracked corresponding differences in decision certainty as measured by the decision’s underlying probability.

### 2.3 Gaze Prediction exhibits statistical hallmarks of confidence

We next examined whether Gaze Prediction exhibits the three statistical hallmarks of subjective confidence proposed by Sanders et al. (2016): (1) a monotonic relation with accuracy, (2) a “folded X” pattern, in which Gaze Prediction increases with increasing trial evidence for correct trials but decreases with increasing trial evidence for incorrect trials, and (3) high-magnitude Gaze Prediction trials show a stronger association between accuracy and evidence than do low-magnitude Gaze Prediction trials (see Fig. 3 and SM for further details of the analysis). Achieving the first hallmark, we found a positive monotonic relation between accuracy and Gaze Prediction, with the group correlation coefficient distribution significantly greater than zero (*Exp. 1*: Mean = .33, 95% CI = [.14, .53], *t*_24_ = 3.5, *p* = .002, Cohen’s *d* =0.7; *Exp. 2*: Mean = .28, 95% CI = [.11, .45], *t*_24_ = 3.4, *p* = .002, Cohen’s *d* =0.61; Fig. 3A and Fig. S7). To test the second hallmark, and following our preregistration plan, we used trial-by-trial expectation strength (*v*) derived from the learning model as a proxy for trial evidence. For correct trials, Gaze Prediction increased with *v,* whereas for incorrect trials Gaze Prediction decreased, exhibiting a “folded X” pattern (see Fig. 3B). The interaction effect of accuracy and *v* was significant (*Exp. 1*: ^2^_(1)_ = 9.32, *p* = .002; Exp. *2*: ^2^_(1)_ = 20.67, *p* < .001; see Eq. 1 in Table 1 and Table S5 and Table S6 for pairwise comparisons). Examining the simple effects of the interaction across both experiments revealed a significant upward slope for correct trials (Mean β = .65, SE = 0.08, ^2^_(1)_ = 70.65, *p* < .001) and a significant downward slope for incorrect trials (Mean β = -.3 , SEM = 0.15, ^2^_(1)_ = 4.10, *p* = .04; see SM for further control analyses that corroborate this result). Next, we examined whether the folded X is dependent on the learning process and the trial’s relative position in the block. The folded X’s interaction effect was significantly greater for trials in the latter part of the block as opposed to its beginning (*F*_2,76_ = 8.96, *p* < .001, η^2^ = 0.13), and this was driven primarily by a more pronounced downward slope for incorrect trials in later parts of the block (*F*_2,76_ = 7.16, *p* = .001, η^2^ = 0.11; see Fig S8 and SM). Thus, as the block proceeds and learning improves, the folded X becomes more pronounced. Finally, fulfilling the third hallmark, high Gaze-Prediction trials exhibited a stronger association between *v* and accuracy than did low Gaze-Prediction trials (see Fig. 2C), and the interaction was significant (*Exp. 1*: Mean (SEM) = 0.15 (0.05), *t*_24_ =3.16, *p* = .002; *Exp. 2*: Mean (SEM) = 0.16 (0.04), *t*_30_ =3.60, *p* < .001; see Eq. 2 in Table 1). Thus, Gaze Prediction exhibits three statistical hallmarks previously observed in reported confidence. We therefore provisionally use the label “gaze-based confidence” when comparing it to explicit confidence judgments in the following sections.

**Figure 3.**
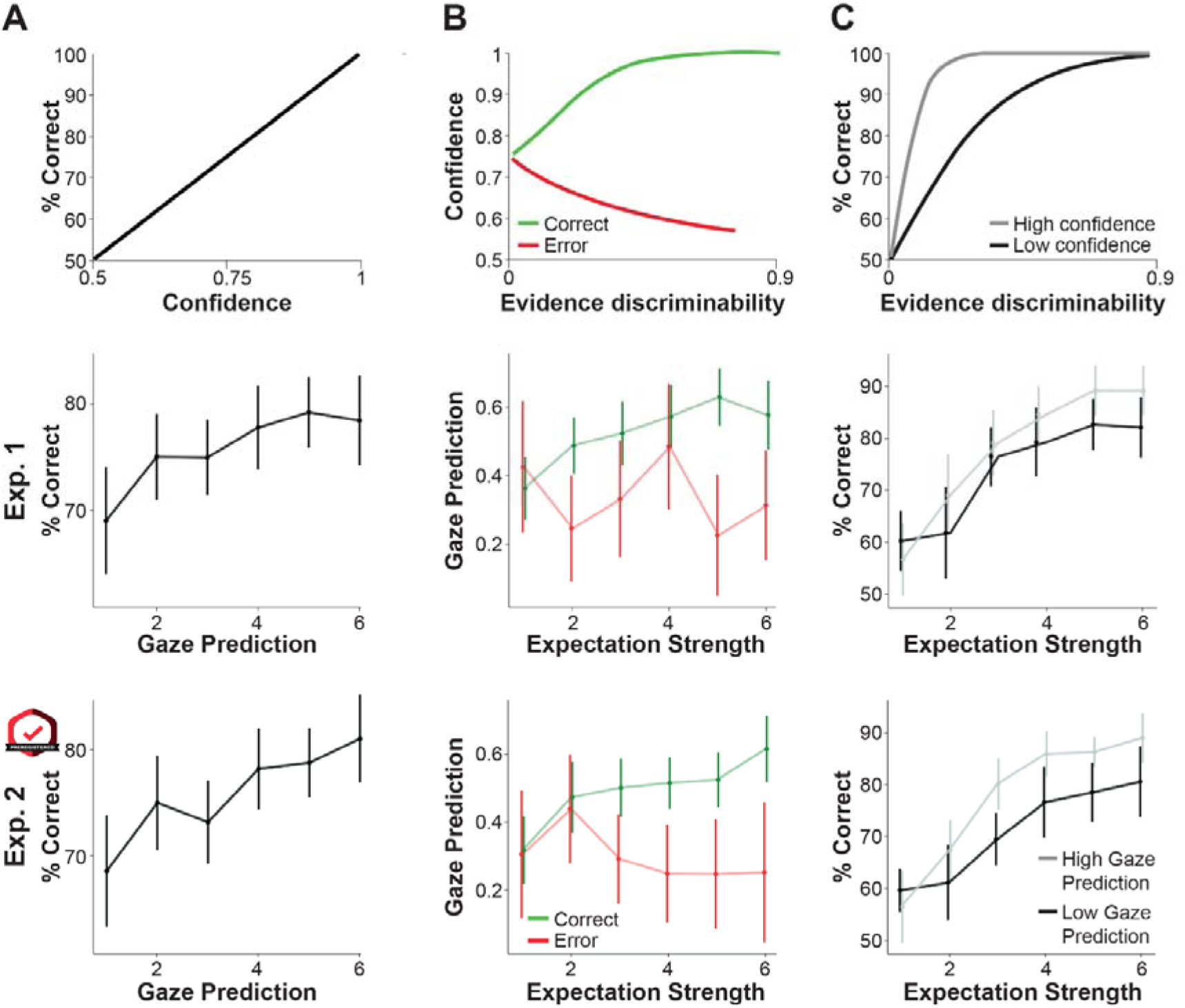
Gaze Prediction exhibits statistical hallmarks of confidence. **Top panel:** Sanders et al.’s (2016) three statistical hallmarks of confidence: (a) A monotonic relation between confidence and accuracy, (b) a “folded X” pattern, such that as evidence increases, confidence in correct trials (green) increases, whereas for incorrect trials (red) it decreases, and (c) steeper slope between evidence strength and accuracy for high confidence trials (gray) compared to low confidence trials (black). Figure adapted from Sanders et al., (2016). **Middle and bottom panels:** Gaze Prediction, the continuous degree of alignment between post-decision gaze direction and predicted target direction, exhibits computational hallmarks of confidence in an exploratory (Experiment 1) and preregistered replication (Experiment 2) experiment. In these analyses, Gaze Prediction was used in lieu of confidence, and expectation strength (*v*) served as a proxy for trial evidence. Circles indicate group mean and error bars are 95% CI.

**Table 1.**
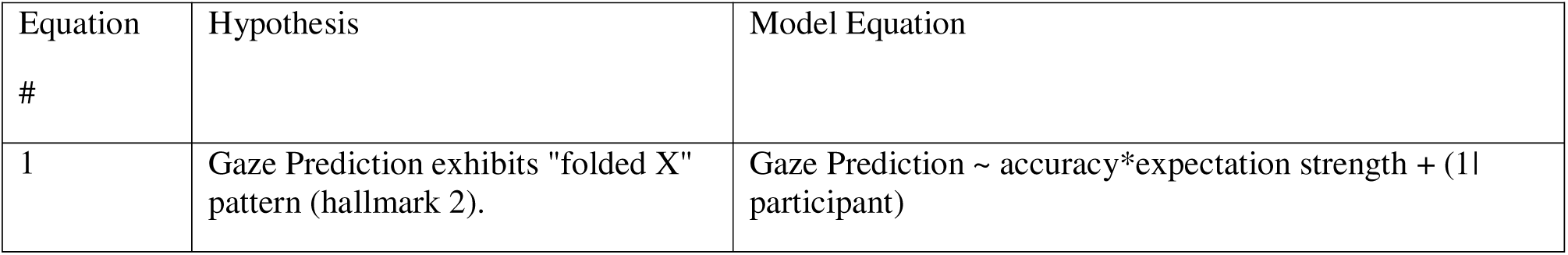
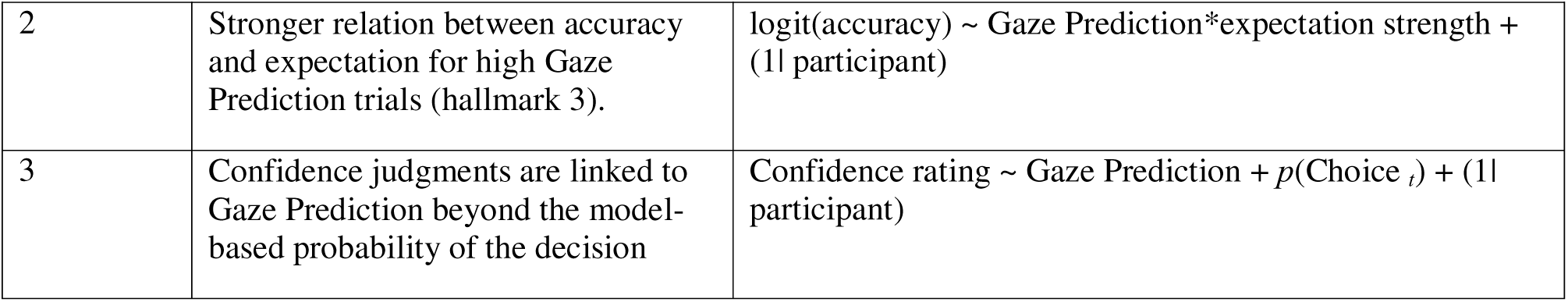
Regression Models.

### 2.4 Gaze-based confidence is significantly but modestly correlated with confidence ratings

In Experiment 3, after predicting the target’s location, participants rated their confidence in their decision (see SM for complete results of the experiment’s preregistered analyses). First we examined the relation between explicit confidence ratings and trial-by-trial variation in decision certainty, indexed by *p*(Choice *_t_*). The Pearson correlation between the two measures was calculated separately for each participant, and the group-level distribution of correlation coefficients was found to be significantly greater than zero (Mean *p*= .38, 95% CI = [.33, .44], *t*_31_ = 14.47, *p* < .001, Cohen’s *d* = 2.56; see Figure S18 for correlations to other learning variables). In addition, the correlation was statistically reliable at the individual level for 30 of the 32 participants.

Turning to examine the relation between confidence ratings and gaze-based confidence, we calculated the trial-by-trial correlation between confidence ratings and Gaze Prediction for each participant. The group-level distribution of correlation coefficients was significantly greater than zero (Mean group *p*= .076, 95% CI = [.038, .11], *t*_31_ = 4.1, *p* < .001, Cohen’s *d* =0.72; see Figure 4A), though the magnitude of the mean trial-level correlation was below our preregistered criterion (*r* >. 1) for defining the two measures as “similar.” Because both confidence ratings and Gaze Prediction were found earlier to correlate with *p*(Choice *_t_*), we next modeled confidence ratings as a function of Gaze Prediction, controlling for *p*(Choice *_t_*). Using a linear mixed-effects model (see Eq. 3), the conditional association between Gaze Prediction and confidence ratings remained significant, albeit with a small effect size (χ²_(1)_ = 11.62, *p* < .001, ΔR^2^ = .002, Cohen’s *f*^2^ =.003). This result was further corroborated by fitting the same model separately for each participant and examining the group-level distribution of the beta coefficients (Mean β _Gaze Prediction_ = 0.07, 95% CI = [.021, .13], *p* = .006, Cohen’s *d* =.52; Mean β *_p_*_(Choice_ *_t_*_)_ = 1.67, 95% CI = [1.42, 1.92], *p* < .001, Cohen’s *d* = 2.38). Finally, a significant conditional association was also observed when controlling for the potential contribution of additional confounding variables (i.e., response accuracy and previous trial’s prediction error) (χ²_(1)_ = 9.5, *p* = .002; see SM for results and corroborating analyses using a group-level approach). Taken together, these results indicate that explicit and gaze-based confidence share a small but reliable variance component that cannot be accounted for by the modeled decision certainty indexed by *p*(Choice *_t_*).

**Figure 4.**
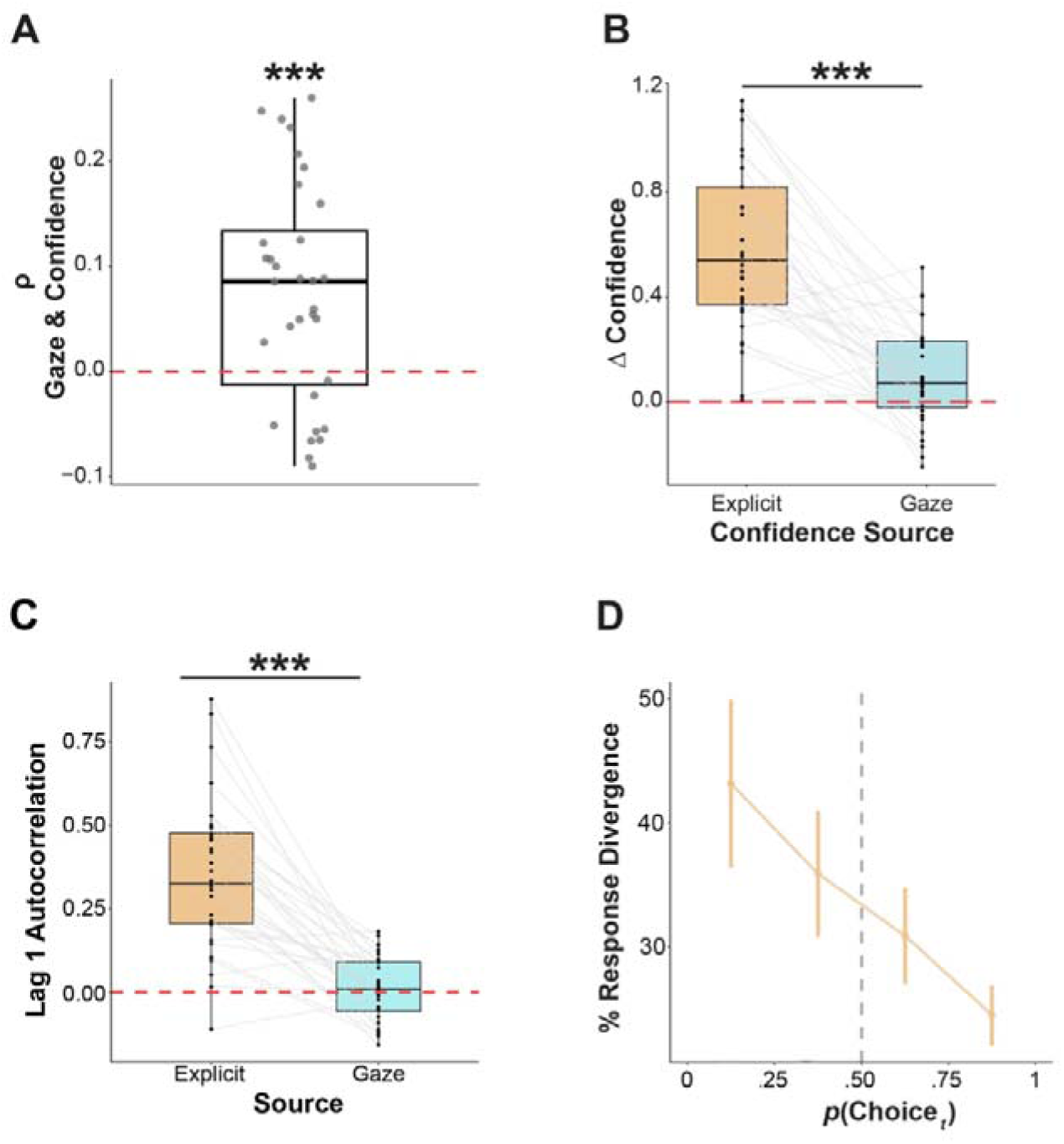
Comparison of reported and gaze-based confidence. **Panel A: Distribution of participants’ correlation coefficients between reported and gaze-based confidence.** Mean correlation was significantly greater than zero (*p* < .001), yet its magnitude was smaller than our preregistered criterion for defining the two processes as similar. **Panel B: Comparison of metacognitive sensitivity of reported and gaze-based confidence.** Reported confidence ratings were significantly more sensitive than gaze-based confidence (*p* < .001). However, the sensitivity of both types of confidence was greater than zero (dashed red line; both *p*s < .001). **Panel C: Comparison of serial dependence in reported and gaze-based confidence.** Reported confidence exhibited significantly greater autocorrelation than gaze-derived confidence (*p* < .001), suggesting a more temporally extended assessment. **Panel D: Increased gaze–decision divergence under decreased decision certainty.** Gaze and decision binary direction diverged more frequently as decision certainty indexed by the model-based decision probability (*p*(Choice *_t_*)) decreased. Exploratory trials *p*(Choice *_t_*) < .5; dashed grey line) diverged more than exploitation trials *p*(Choice *_t_*) > .5. Error bars are 95% CI.

### 2.5 Explicit confidence exhibits superior metacognitive sensitivity but also greater serial dependence compared to gaze-based confidence

After establishing the partial overlap of explicit and gaze-based confidence, we elucidate their common and distinct functional characteristics. For comparability, Gaze Prediction was converted into a 5-point rating analogous to the confidence ratings (see SM for further details). We compared the metacognitive sensitivity of rated and gaze-based confidence in terms of the difference in confidence between correct and incorrect trials (Δ confidence; see SM for justification of this measure). Both measures exhibited significant metacognitive sensitivity, with higher confidence in correct trials compared to incorrect trials (i.e., Δ Confidence > 0; see Figure 4B, Δ Confidence *_Reported_*: Mean = 0.58, 95% CI = [.47, .70], *t*_31_ = 10.44, *p* < .001, Cohen’s *d* =1.85, Δ Confidence *_Gaze-based_*: Mean = 0.10, 95% CI = [.03, .17], *t*_31_ = 3.01, *p* = .004, Cohen’s *d* =0.54). However, confidence ratings exhibited significantly greater sensitivity than gaze-based confidence (*t*_31_ = 7.7, *p* < .001, Cohen’s *d* = 1.36). Interestingly, metacognitive sensitivity for reported and gaze-based confidence was not significantly correlated across participants, and Bayesian statistics yielded anecdotal evidence for the lack of correlation (*r_31_* = .09, *p* = .63, BF_10_ = .43; see Fig. S9) that should be interpreted with caution given the small sample size.

To further explore possible differences in the mechanism behind each type of confidence, we examined whether they are influenced by the previous trial’s confidence level. To measure serial dependence, for each participant, we calculated the autocorrelation between adjacent trials’ confidence (i.e., lag 1) separately for reported and gaze-based confidence. Reported confidence ratings were autocorrelated across trials, and the group distribution of the coefficient was significantly greater than zero (*M* _autocorrelation_ = 0.35, 95% CI = [.27, .43], *t*_31_ = 8.77, *p* < .001, Cohen’s *d* =1.54). In contrast, the mean group-level autocorrelation of gaze-based confidence was less robust: In Experiments 1 and 2 (combined) it was significantly greater than zero (*M* _autocorrelation_ = 0.05, 95% CI = [.02, .07], *t*_55_ = 3.40, *p* = .001, Cohen’s *d* =0.45), but in Experiment 3 it was not (*M* _autocorrelation_ = 0.01, 95% CI = [-.02, .05], *t*_31_ = 0.74, *p* = .46). Furthermore, directly comparing the autocorrelation coefficient of reported and gaze-based confidence for each participant in Experiment 3 revealed that reported confidence was more strongly autocorrelated than gaze-based confidence (see Figure 4C, *t*_31_ = 7.85, *p* < .001, Cohen’s *d* = 1.39; see SM for similar results when controlling for additional factors such as preceding-trial accuracy). Thus, whereas reported confidence is characterized by a substantial degree of serial dependence across adjacent trials, gaze-based confidence is more trial-specific.

### 2.6 Gaze diverges from the predicted direction under decreased decision certainty

As a behavioral manifestation of internal confidence, gaze—decision alignment may directly contribute to adaptive behavior. To explore this idea, we analyzed the convergence and divergence between the predicted target direction and the binary post-decision gaze direction (Right or Left). As previously noted, post-decision gaze was predominantly directed toward the predicted location in approximately 70% of the trials. The remaining, diverging trials were characterized by a lower level of decision certainty, as indexed by *p*(Choice *_t_*). Exploratory trials, defined as those in which *p*(Choice _t_) < .5, exhibited a significantly higher rate of gaze–prediction divergence than exploitation trials (*p*(Choice *_t_*) >.5; see Figure 4D, *M* _divergence in exploratory trials_: 35%, 95% CI = [31%, 40%]; *M* _divergence in exploitation trials_: 26%, 95% CI = [23%, 28%]; *t* = 4.13, *p* = .001, Cohen’s *d* = 0.56; see SM and Fig. S10). Accuracy was also significantly lower in diverging trials (*M* _accuracy in diverging trials_: 71%, 95% CI = [68%, 74%]; *M* _accuracy in converging trials_: 78%, 95% CI = [76%, 79%]; *t* = 4.75, *p* < .001, *d* = 0.63; see SM Fig. S11). Thus, the predominant gaze direction diverged from the predicted location more often when the decision certainty, as measured by its model-based probability, was relatively low. These results suggest that gaze—decision alignment may also provide a “hedging” mechanism, by which post-decision gaze is directed in the opposing direction at a rate that is inversely related to the decision probability.

## 3. Discussion

This study investigated how post-decision gaze behavior relates to confidence in visual-spatial decisions during probabilistic learning. As expected, across three experiments, the overall direction of the participants’ gaze aligned with the immediately preceding left– right prediction, matching the predicted direction on roughly 70% of the trials. More importantly, in all three experiments, the degree of gaze–prediction alignment tracked the underlying model-based probability of the decision (i.e., *p*(Choice)), a computational index of decision certainty, related to second-order processes during learning (Hertz et al., 2018; Meyniel et al., 2015). In addition, gaze–prediction alignment exhibited the three statistical hallmarks previously observed in reported confidence (Sanders et al., 2016). Finally, directly probing the relation between gaze–prediction alignment and explicit confidence ratings in Experiment 3 yielded a significant trial-level correlation even when controlling for the model-based decision probability that indexes decision certainty. Together, these results demonstrate that post-decision gaze behavior exhibits multiple confidence-like characteristics linked to decision certainty. Thus, in the present visual-spatial paradigm, the strength of gaze–prediction alignment appears to provide a behavioral index of internal confidence during probabilistic learning.

The sensitivity of post-decision gaze to decision certainty likely serves a functional role. It may be part of an anticipatory motor preparation process, directing attention toward the predicted location to facilitate reaching. While strong commitment facilitates movement when correct, it is detrimental if the prediction is wrong. Thus, gaze–prediction alignment represents the modulation of instrumental behavior. A complementary mechanism may be representational. Post-decision gaze could reflect the continued activation of internal models (Viganò et al., 2024). Stimulus activation in working memory often triggers involuntary gaze shifts to external locations, even when task-irrelevant (van Ede et al., 2020). Here, active representations of the predicted target and its reliability may automatically bias gaze in the predicted direction.

Post-decision gaze exhibited statistical hallmarks previously observed in confidence (Sanders et al., 2016), most notably the folded-X pattern. This pattern is likely driven by two complementary processes. First, the downward slope for incorrect trials may reflect strategic “hedging”. When decision certainty is low, participants are more likely to divert their gaze towards the other option. Accordingly, the folded-X pattern became more pronounced later in each block, when learning improved and expectations were stronger. Under these conditions, occasional exploratory choices, that are characterized by lower decision probability reflecting increased uncertainty, are more likely to be associated with reduced gaze-decision alignment. Second, the pattern may reflect dynamic post-decisional evidence integration, similar to those observed in perceptual tasks (Moran et al., 2015; Van den Berg et al., 2016). Extensive work in value-based decision-making shows that perceived value is not static but evolves dynamically within a trial, often captured by accumulator models (Krajbich et al., 2010). Importantly, confidence ratings closely track these evolving value fluctuations within trials (De Martino et al., 2013). Future work should examine the extent to which these processes overlap with or diverge from mechanisms underlying confidence ratings folded-X pattern during probabilistic learning. Importantly, the extent to which these hallmarks, and in particular the folded-X pattern, warrant the interpretation of post-decision gaze as confidence-driven sensorimotor behavior echoes similar debates regarding other measures, such as neural activation and animal behavior that exhibit these hallmarks (Bang & Fleming, 2018; Rausch & Zehetleitner, 2019; Schmack et al., 2021).

When interpreting these findings as supporting the contribution of second-order processes to post-decision gaze, it is important to also consider the possibility that gaze might solely reflect probabilistic expectations of target location. Clearly, gaze behavior, both in the present task and in general, primarily reflects a probabilistic expectation of the target’s location. However, does post-decision gaze also include a component specifically related to internal confidence? Empirically distinguishing between these accounts in the current paradigm is challenging because the model-based decision probability can support both interpretations. Post-decision gaze consistently tracked the model-based decision probability, a measure that has been previously linked to decision confidence in probabilistic learning (Hertz et al., 2018; Meyniel et al., 2015). In addition, gaze was also significantly correlated with explicit confidence ratings in Experiment 3. Critically, Experiment 3 provided initial support for a second-order component of gaze, by demonstrating that gaze-decision alignment and explicit confidence ratings were significantly correlated even after controlling for model-based decision probability. Although the residual relationship was small in magnitude, its existence is not trivial, given that second-order evaluations are also partially driven by decision certainty as indexed by decision probability (Fleming & Daw, 2017; Maniscalco & Lau, 2012). Further possible support for a second-order component of post-decision gaze can be found in the task’s structural timing. Committing to an explicit prediction may trigger a transition from evidence accumulation to performance monitoring and maintaining an active representation of the pre-gaze prediction in working memory during the post-decisional interval to guide gaze dynamics (Ben Yehuda et al., 2025; Meyniel et al., 2015; Pouget et al., 2016; Yeung & Summerfield, 2012). These theoretical considerations and empirical findings provide initial support for the possibility that post-decision gaze is modulated by both the underlying decision probability and second-order processes. Importantly, gaze-based confidence and explicit confidence judgments should not be conceptualized as equal manifestations of internal confidence. Unlike reported confidence, which involve a deliberate evaluation and report of one’s internal confidence, gaze-based confidence reflects the modulation of world-directed instrumental behavior by second order influences and is thereby an indirect expression of confidence.

The different characteristics of explicit confidence ratings and gaze-based confidence likely reflect their distinct functional roles. First, consistent with previous work (Mei et al., 2023; Rahnev et al., 2015, 2020), confidence judgments showed significant serial dependence. In contrast, gaze-based confidence showed substantially weaker and less robust serial dependence. Serial dependence in confidence judgments has been proposed to reflect a metacognitive “continuity field”, that is a temporally extended assessment of decision performance that persists across trials (Rahnev et al., 2015). In probabilistic learning tasks, in which successive trials are probabilistically contingent, some degree of serial dependence may contribute to a more coherent and stable (and perhaps more accurate) assessment of overall performance. Thus, reported confidence may reflect the contribution of a higher-order metacognitive decision process that enhances self-consistency over time (Koriat, 2012). In contrast, gaze-based confidence and other implicit behavioral measures may involve a more direct and less stabilized influence of an internal confidence signal. A second difference between gaze-based and reported confidence is their relative level of metacognitive sensitivity. Although the gaze-based measure showed significant sensitivity, it was substantially lower than for reported confidence. This likely reflects that, unlike explicit reports, the primary function of post-decision gaze behavior is not to provide an accurate index of internal confidence. Instead, post-decision gaze is primarily instrumental, with confidence being only one of many factors influencing its dynamics. Taken together, these differing characteristics of partially overlapping manifestations of confidence in the same decision may be shaped by their distinct functional roles.

Understanding how functionally differentiated explicit and implicit manifestations of internal confidence are coordinated within a unified behavioral system is important for characterizing adaptive decision-making and behavior. This may be relevant to the understanding of functional impairment in psychopathologies, such as schizophrenia spectrum disorders, in which aberrant interaction between explicit and implicit-sensorimotor systems is postulated to be a core deficit (Krugwasser et al., 2021; Salomon et al., 2021; Stern et al., 2020).

## 4. Materials and Methods

We conducted three experiments: an initial exploratory Experiment 1, and two preregistered experiments (Experiment 2: https://osf.io/p4hyq, 09.05.2023; Experiment 3: https://osf.io/ngsx6, 17.07.2023). The preregistered analysis plan was followed, unless otherwise stated. For consistency, we retrospectively executed the preregistered analysis plan for Experiment 1. The primary inclusion criteria were: (1) actual accuracy (i.e., the prediction matches the target’s actual location) above 55%. This value was selected to ensure that parameter fitting was reliable and that participants understood the task; (2) The VR and eye-tracking recording was valid in at least 70% of trials.

### 4.1 Participants

For Experiment 1 we collected data from 35 participants; 25 participants (19 female, *M*_age_ = 23.9, *SD* = 4 years) were included in analysis. For Experiment 2, we preregistered 40 participants, anticipating that ∼30 would provide sufficient statistical power to replicate Experiment 1’s effects (see preregistration for power analysis). The final sample included data from 31 participants (21 female, *M*_age_ = 23.7, *SD* = 3.4 years). For Experiment 3, we preregistered 40 participants, and the final sample size included data from 32 participants (23 female, *M*_age_ = 25.3, *SD* = 5.3 years).

All participants were right-handed, with normal or corrected-to-normal vision and self-reported no history of neurological impairments and were recruited via social media. All participants gave written informed consent and were compensated (monetary compensation or course credits) for their participation. Experiment 1 and 2 were performed at Bar-Ilan University and Experiment 3 was performed at Haifa University. Each university’s Institutional Review Board approved this study: Approval ISU202107003 for Experiments 1 and 2 and Approval HU448/19 for Experiment 3.

### 4.2 Hardware and Setup

Participants wore an HTC Vive Pro Eye head-mounted display connected to a high-performance desktop, which projected a VR environment created in Unity (2018.3.2) at a 90 Hz refresh rate and tracked their hand and eye movements. Responses were recorded through the controller’s trackpad, and eye movements were monitored using the built-in Tobii trackers (∼90Hz; see SM for additional technical details).

### 4.3 Experimental Procedure

We adapted a well-established previously modeled probabilistic learning task (Harrison et al., 2021; Iglesias et al., 2013) to VR. In 160 trials, participants were presented with one of two probabilistic spatial cues (i.e., a square that appeared in one of two colors), and then predicted the upcoming target’s location (Right/Left) by pressing a button on the controller. Following their prediction, a fixation cross appeared (duration 1.5 seconds), and then participants entered a virtual environment and the target butterfly appeared after a brief 300 millisecond delay. The butterfly appeared equally often in one of six possible locations, distributed symmetrically across the left and right sides of the environment. While the target remained at a static coordinate it performed a continuous wing-flapping animation. Participants were instructed to reach towards the target as quickly and accurately as possible once they identified it, and a virtual hand tracked their actual hand movement (see Fig. 1A for trial flow).

The participants’ goal was to learn the cue-location mapping rule underlying a series (block) of trials and make their predictions accordingly. Participants were informed that: (1) For each trial, one of two opposing mapping rules was in effect: Under one rule, cue A indicated a leftward target and cue B indicated a rightward target; under the alternative rule, these mappings were reversed. (2) These cue–location mappings were probabilistic, such that in 75% of trials the target appeared in the indicated location (valid-cue trials), whereas in the remaining 25% it appeared in the opposite location (invalid-cue trials). (3) Over the course of the experiment, the rule governing the current series of trials changed multiple times, requiring participants to detect these changes and adjust their predictions accordingly. Unbeknown to the participants, the rule was changed five times during the experiment, with each block lasting between 29 and 35 trials (see SM for further details). In Experiment 3 participants performed an identical task and additionally rated their confidence in their prediction on a five-point vertical sliding scale. After completing the probabilistic learning task, participants also performed a sensorimotor conflict-detection task (Applebaum et al., 2025) and completed self-report questionnaires for exploratory purposes (see SM).

## 5. Measures

In each trial, we recorded participants’ prediction (“The target will appear in the Right or Left location”) and their post-decision ocular behavior. Predictions were assessed for *actual accuracy* (the target appeared as predicted) and *rule accuracy* (the prediction conformed to the underlying rule). Our primary focus was on rule accuracy due to our interest in the rule-learning process, except when noted otherwise.

Gaze direction was measured across a preregistered 300 millisecond interval, beginning upon entry into the VR environment (1500 millisecond after making the left-right prediction) and ending with the onset of the target butterfly. To account for individual variability, horizontal gaze eccentricity was first normalized (with 0 corresponding to the viewer-centered midpoint) across all trials for each participant. A time-weighted average of these normalized values was then calculated for each trial across the 300-millisecond interval, yielding a single continuous Gaze Direction measure: a composite measure of the duration and magnitude of lateral gaze bias, with positive and negative values representing the strength of rightward and leftward bias, respectively. From this measure, we derived two additional measures: (1) a binary measure analogous to the response made via the controller, based on the sign of the Gaze Direction measure and indicating the predominant gaze direction; and (2) *Gaze Prediction*—a continuous measure of gaze–prediction alignment, derived by adjusting the sign of Gaze Direction to indicate convergence with (positive sign) or divergence from (negative sign) the predicted target direction. Conceptually, this measure indexes the graded degree of alignment between post-decision gaze direction and the preceding binary decision on each trial, with positive values scaling the degree of alignment in the predicted direction, and negative values scaling the degree of misalignment in the opposite, diverging direction (see SM for transformation details).

### 5.1 Modeling

To analyze learning dynamics throughout the experiment, we adopted a model-based approach, using the well-established family of Rescorla-Wagner (RW) reinforcement-learning models ( (Rescorla, 1972); see SM for further details of the modeling process). Deviating from our preregistered plan to use a simple RW model, after conducting formal model comparisons we chose to use a Rescorla-Wagner model with an additional stickiness parameter (sRW; Culbreth et al., 2016; Daw et al., 2011) given its superior fit (see SM).

In brief, the sRW model consists of a perceptual and decision model. In the perceptual model, each option’s value (*v*) is updated by the prediction error (δ)—the discrepancy between the previous trial’s predicted and actual outcomes (*r*)—modulated by the learning rate (), a free parameter that controls the extent of updating by prediction errors (Eq. 4A). Lower values reflect slower learning over a larger number of trials. The value (*v*) from the perceptual model informs the decision model, which employs a SoftMax function to calculate the choice probability, *p*(Choice *_t_*). The function’s steepness and randomness are dictated by another free parameter, inverse temperature (β), reflecting decision consistency (Eq.4B). It also includes a perseveration (“stickiness”) parameter (⍰) to quantify the tendency to use the rule used in the previous decision, multiplied by *rep (a)*, an indicator function that equals 1 for decisions that apply the rule used in the previous decision and 0 otherwise.

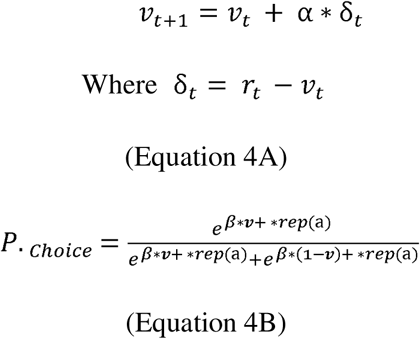

In line with our preregistration and previous work, stimuli and responses were encoded within a unified contingency space to minimize free parameter estimates (Harrison et al., 2021; Iglesias et al., 2013). Model parameters were estimated via the Bayesian adaptive direct search (BADS) toolbox (Acerbi & Ji, 2017), optimizing each model’s log likelihood. Parameter recovery across models was reliable (see Fig. S17). Model comparison was performed via the Bayesian Information Criteria (BIC; (Schwarz, 1978)). All models showed good model recovery (see Fig. S18).

Linear mixed models were fitted using the lme4 package (Bates et al., 2015), and likelihood ratio tests were used to compare the full model against a version without the targeted effect to obtain p-values.

## Data and materials availability

Pre-registrations, analysis code and notebooks available at https://osf.io/qtg6k/.

Data will be made public immediately following publication, currently available for reviewers at: https://osf.io/6nj8u/?view_only=0852199aacb94825a3ef81c760b972a6

## Author contributions

Conceptualization: YS, ON, DK, MG, UH, RS

Methodology: YS, ON, UH, YZ, RS

Investigation: YS, ON, UH, RS

Visualization: YS, UH, RS

Supervision: UH, DK, RS

Writing—original draft: YS, UH, RS

Writing—review & editing: YS, ON, MG, DK, YZ, UH, RS

