## Supplemental Material for "Post-Decision Gaze as a Behavioral Manifestation of Decision Confidence"

for

### Results

#### Decisions reflect learning of probabilistic rule across experiments

We examine whether participants’s decisions exhibit signatures of probabilistic learning at the group-level across experiments. We examine here, across all participants, three well established model-free signatures of probabilistic learning, that also served as pre-registered inclusion criteria. First, overall actual accuracy (i.e., the predicted location of the target matched its actual location) was significantly greater than chance (see Figure S1, Table S1) and was also greater than the pre-registered criteria of 55%. Second, accuracy on trials in which the cue was valid was greater than trials in which the cue was invalid (see Figure S1, Table S2). Third, accuracy within a block, (i.e., consecutive trials governed by the same underlying rule), exhibited the expected learning curve. This was true both for actual accuracy and rule accuracy (see Figure S2).

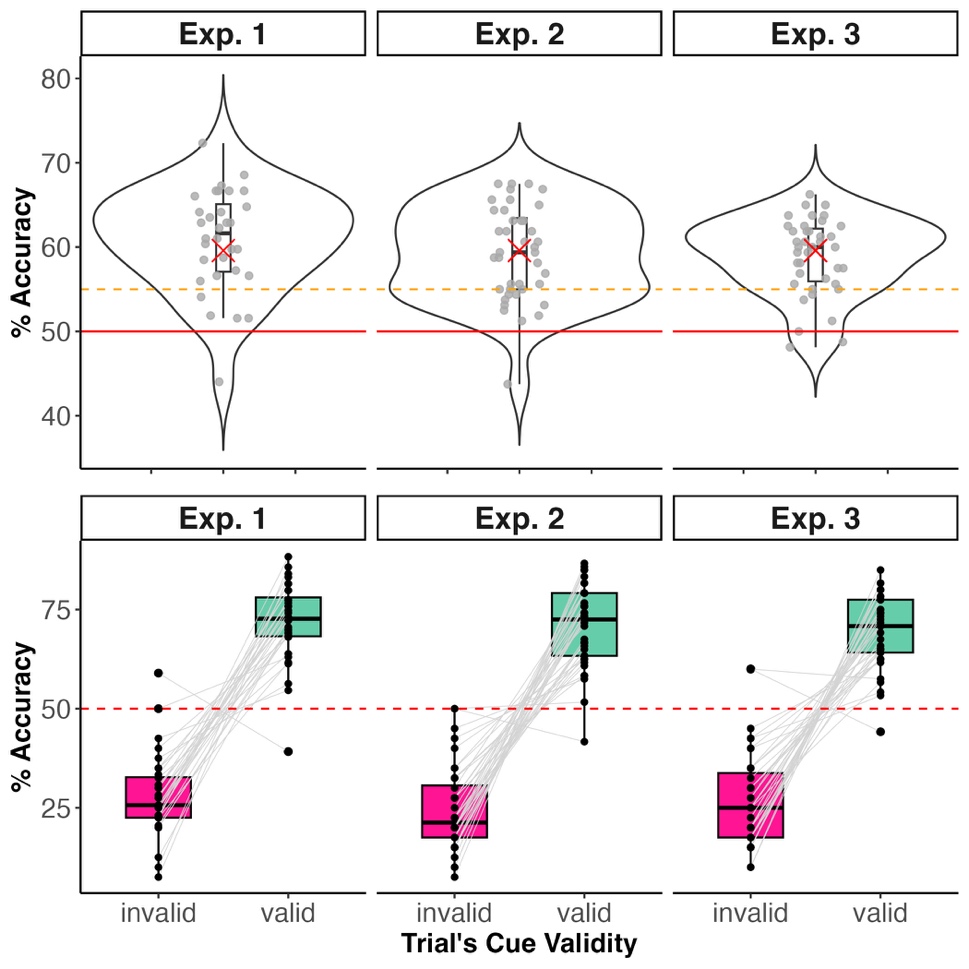

**Figure S1. Signatures of probabilistic learning across all experiments and all participants. (Top panel)** Overall actual accuracy per experiment. Accuracy here refers to whether the explicit prediction matched the target’s location. Solid red line is chance performance, and dashed orange line is pre-registered 55% accuracy inclusion criteria. **(Bottom panel)** Accuracy on trials with valid and invalid cues. Following the pre-registration, participants that did not exhibit higher accuracy on trials with valid cue as opposed to invalid, were not included in main analyses.

| Experiment | Mean Acc. [95% CI] | t_df_ | *p* value | Cohen’s *d* |
| --- | --- | --- | --- | --- |
| Exp. 1 | 60.8 [58.7, 63.0] | t_31_ = 10.2 | *p* < .001 | 10.1 |
| Exp. 2 | 59.2 [57.4, 61.0] | t_39_ = 10.4 | *p* < .001 | 10.6 |
| Exp. 3 | 58.9 [57.5, 60.4] | t_38_ = 12.3 | *p* < .001 | 13.0 |
| Combined | 59.6 [58.6, 60.6] | t_110_ = 18.8 | *p <* .001 | 11.1 |

**Table S1. Accuracy of all participants per experiment.** A one-sample t-test against a mean of 50 was performed, robustly supporting that group performance was greater than chance.

| Experiment | Mean difference [95% CI ] | t_df_ | *p* value | Cohen’s *d* |
| --- | --- | --- | --- | --- |
| Exp. 1 | 44.0 [37.0, 50.9] | t_31_ =12.9 | *p* < .001 | 2.3 |
| Exp. 2 | 46.4 [40.1, 52.7] | t_39 =_ 14.8 | *p* < .001 | 2.3 |
| Exp. 3 | 43.3 [36.6, 50.0] | t_38_ = 13.1 | *p* < .001 | 2.1 |
| combined | 44.6 [40.9, 48.3] | t_110_ = 23.7 | *p <* .001 | 2.3 |

**Table S2. Difference in accuracy on valid and invalid cue trials**. As expected, a paired t-test robustly exhibited increased accuracy on valid trials, demonstrating that participants learned the underlying rule.

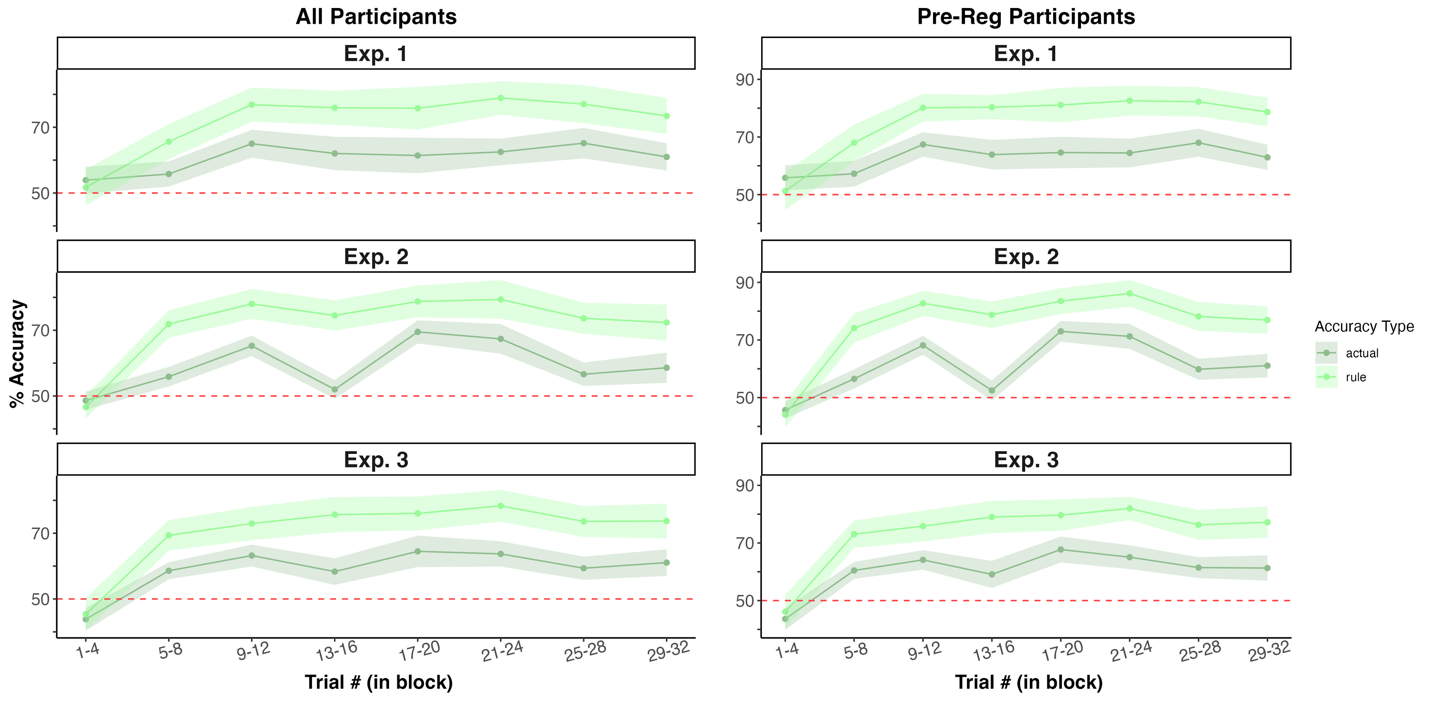

**Figure S2. Learning curve within block across experiments.** Accuracy within blocks of trials (i.e., consecutive trials governed by the same rule) was averaged for each participant across blocks. Trials were binned by their relative position. The shaded area represents the 95% confidence interval across participants. Both 'actual' accuracy (prediction matched target’s actual location) and 'rule' accuracy (prediction matched the underlying rule’s target location) showed the expected learning curve. Rule accuracy was consistently higher than actual accuracy presumably due to invalid trials. In experiment 2, where the trial sequence was identical across participants (SM Methods), a dip in actual accuracy occurred mid-block due to the specific trial sequence, but rule accuracy remained stable, indicating participants learned the underlying rule.

#### Gaze direction robustly converges with explicit predictions

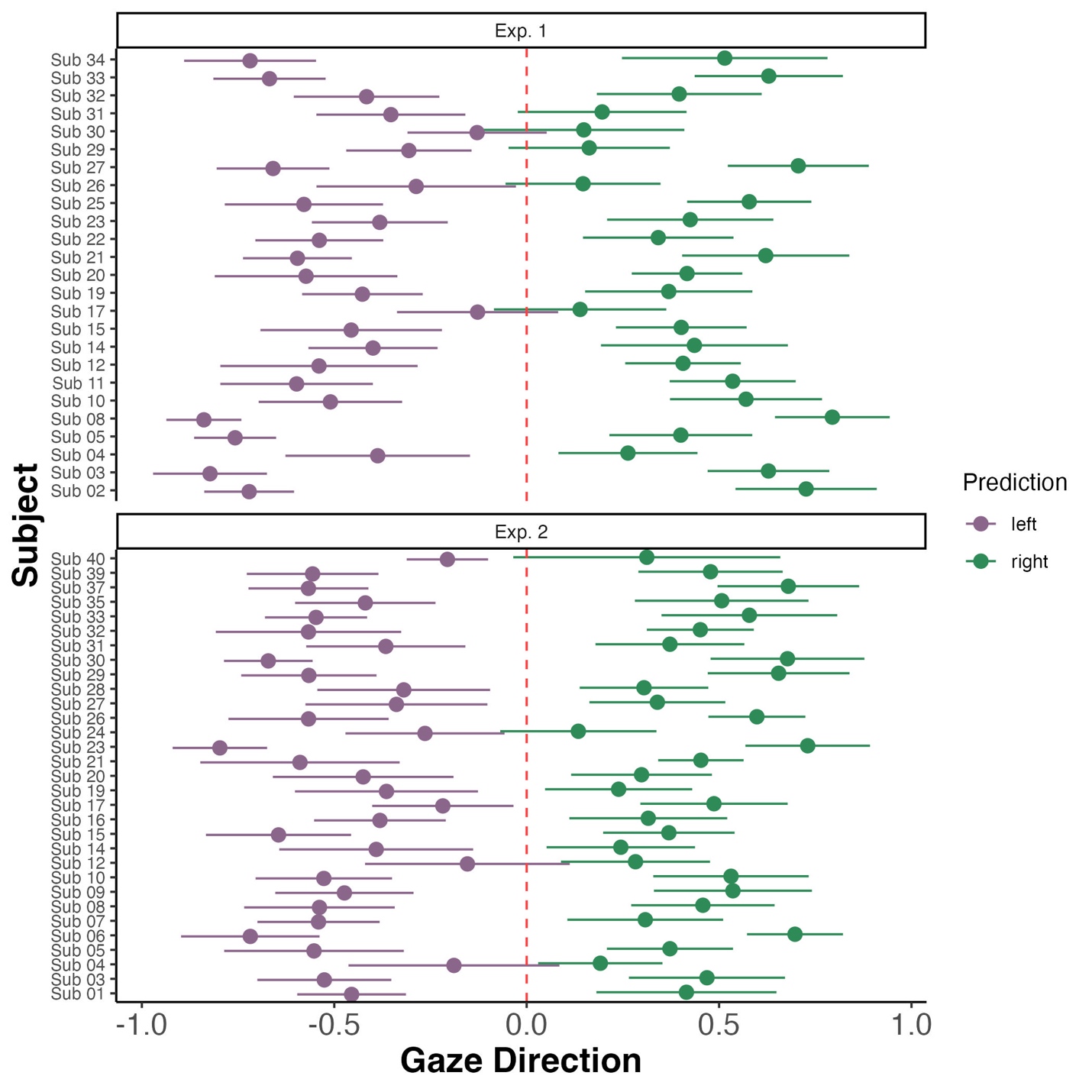

**Figure S3. Gaze direction follows explicit prediction in individual participants.** Mean normalized gaze direction averaged across 300 msec interval prior to target’s appearance is robustly aligned with explicit prediction’s direction (‘right’/ ’left’). Gaze direction was averaged per participant by their explicit prediction, with bars reflecting 95% confidence intervals. Supporting the robust convergence of explicit and ocular predictions, across all participants we see that gaze direction follows the explicit prediction. Furthermore, the divergence was statistically significant at the individual participant level in all participants except for two (Exp. 1’s Sub 30 & Sub 17).

##### Proportion of converging and diverging explicit and ocular responses

Following the high proportion of converging responses reported in the main text and in Figure 2B, here we report the results for the individual experiments that consistently showed similar proportions across the three experiments (see Table S3 & Figure S4).

| Experiment | Explicit and ocular response | Mean proportion [95% CI] |
| --- | --- | --- |
| Exp. 1 | diverge | 26.03 [21.96, 30.09] |
| Exp. 1 | converge | 73.97 [69.91, 78.04] |
| Exp. 2 | diverge | 28.36 [25.20, 31.52] |
| Exp. 2 | converge | 71.64 [68.48, 74.80] |
| Exp. 3 | diverge | 27.23 [23.14, 31.31] |
| Exp. 3 | converge | 72.77 [68.69, 76.86] |

**Table S3. Convergence and divergence of explicit and ocular ‘responses’ across individual experiments.**

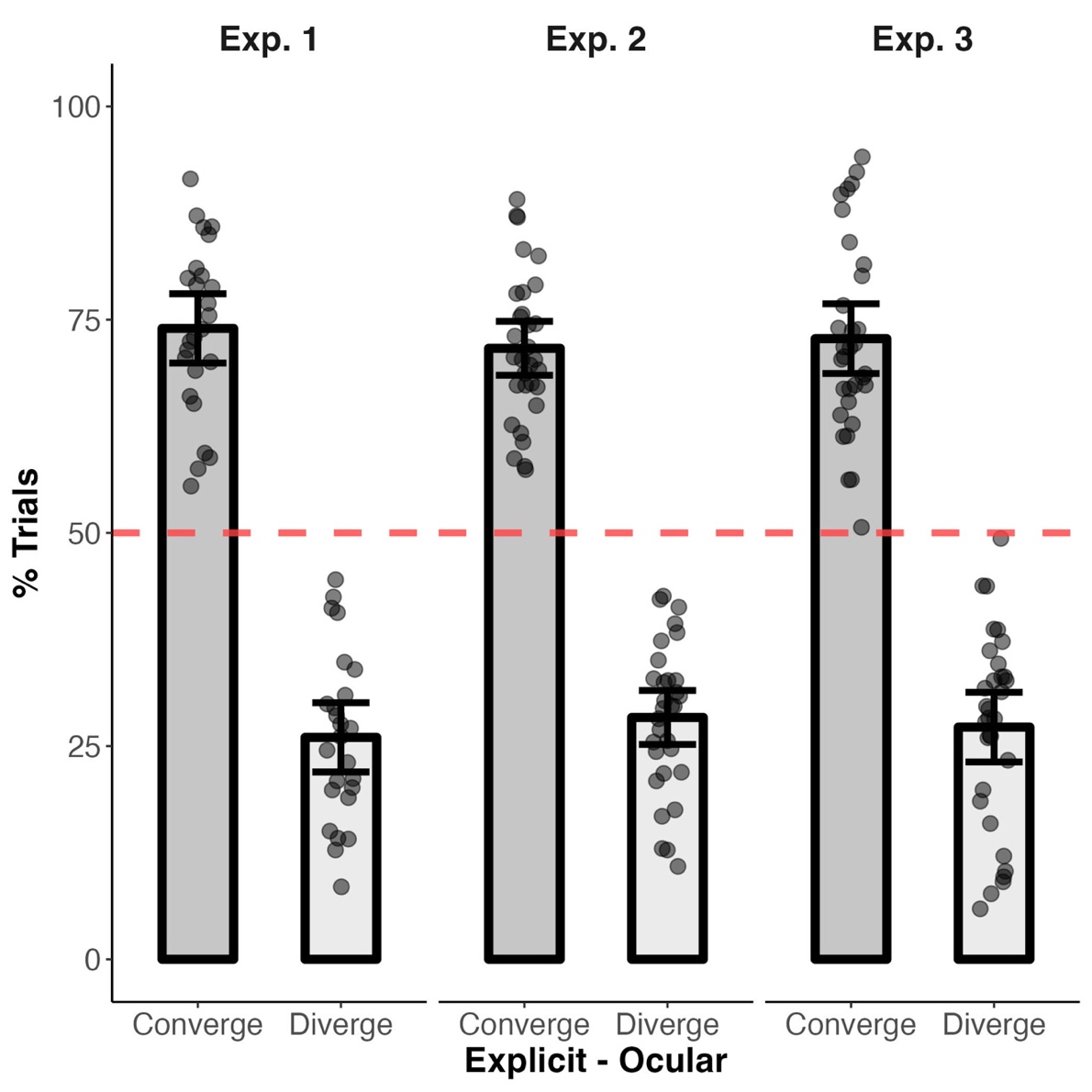

**Figure S4. Convergence and divergence of explicit and ocular ‘responses’ across individual experiments.**

#### Model comparison of explicit and ocular learning

As noted in the Methods, the model space consisted of: (1) the well-established Rescorla-Wagner (RW) model (Rescorla, 1972), (2) a Rescorla-Wagner model with an additional stickiness parameter (sRW) (Culbreth et al., 2016; Daw et al., 2011), (3) and a noisy Win-Stay Lose-Switch (WSLS) model (Wilson & Collins, 2019). Model comparison was performed using Bayesian Information Criteria (BIC; (Schwarz, 1978)) metric.

For explicit decisions the winning model was the sRW, with the lowest BIC (see Table4 & Figure S5) by a difference of ~ 7, reflecting a strong effect (Raftery, 1995). Likewise, a one-way repeated measures ANOVA of BIC values found a significant effect for the model (*Model*: F_1.24,68.02_*= 112.21*, *p* < .001, η^2^*_p_ =* .67).

| **Source** | **Model** | **BIC mean [95 CI%]** |
| --- | --- | --- |
| Explicit | RW | 143.73 [133.51, 153.95] |
| Explicit | sRW | 136.09 [125.03, 147.16] |
| Explicit | WSLS | 194.49 [190.01, 198.98] |

**Table S4. Summary statistics of the model comparison metric BIC for learning models across experiment 1 & 2.**

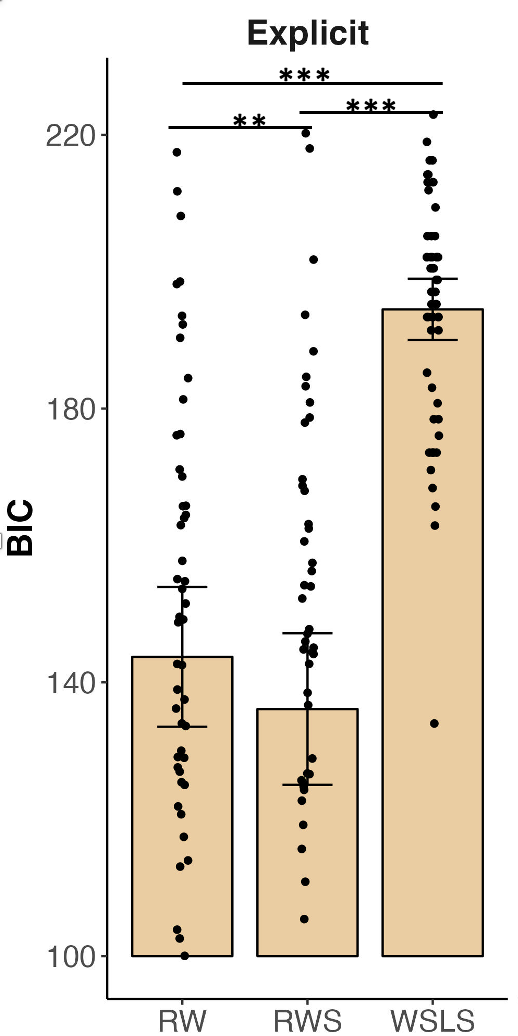

**Figure S5. Model comparison of learning.** Models were compared using BIC. Importantly, lower BIC values indicate a better fit. For the sake of visualization’s clarity, only values above 100 are displayed. For explicit responses the winning model is Rescorla Wagner with a stickiness parameter (RWS). All p values were Bonferroni corrected, ** = *p* < .01; *** = *p* < .001.

Following sRW’s superior fit for explicit responses, we depart from our pre-registered plan and utilize it instead of RW for analyses in which model parameters are obtained from explicit responses in the main text. Importantly, similar results were obtained using the pre-registered RW model.

**Pre-registered results of gaze and explicit prediction convergence using Rescorla Wagner model**

We pre-registered the use of the Rescorla-Wagner (RW) model to investigate latent learning processes and their relationship to ocular expectations. However, our exploratory analyses showed that a modified RW model with a stickiness parameter, the 'sticky RW (sRW) model,' better captures explicit prediction learning (see Fig 5b & Fig S5). Therefore, we used the sRW model in the main text, deviating from our pre-registration.

Here, we present results using the pre-registered RW model. Crucially, all results align with our pre-registered hypotheses, closely resemble those obtained with the sRW model, and are statistically significant.

##### Correlation of *p*(Choice *_t_*) and Gaze Prediction magnitude

We examined whether fluctuations in Gaze Predictions’ magnitude track the probability of the choice made (i.e., *p*(Choice *_t_*)) derived from the RW model. Pearson correlations coefficients were calculated per participant, and the group level distribution was examined. In line with our pre-registration, the group distribution was significantly greater than zero in both experiments (*Exp. 1*: Mean 𝝆 = .15, 95% CI = [.09, .21], *t*_24_ = 5.35, *p* < .001, Cohen’s *d* =1.07, *Exp. 2*: Mean 𝝆 = .13, 95% CI = [.08, .19], *t*_24_ = 5.06, *p* < .001, Cohen’s *d* =0.91; Fig. S6 ).

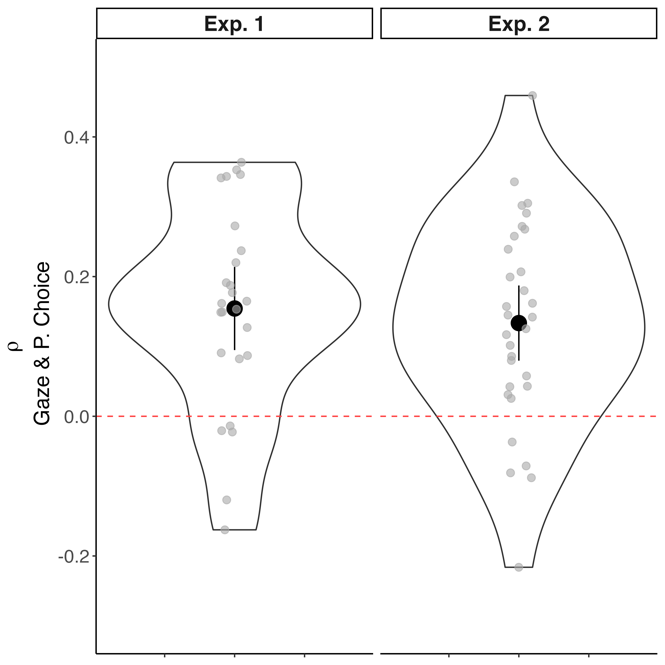

**Figure S6. Distribution of Correlation coefficients between Gaze Prediction and *P. _Choice_* derived from RW model.**

##### Computational Hallmarks of confidence using RW model

We report the results of hallmark 2 and 3 that rely on the trial’s expectation strength (*v*) derived from the computational model using the pre-registered RW model.

In line with the second hallmark, we observed a ‘folded x’ pattern. In correct trials, Gaze Prediction increased with *v,* whereas for incorrect trials’ gaze decreased (see Fig. S7), and the interaction of accuracy and expectation strength was significant (*Exp. 1*: 𝝌^2^_(1)_ = 11.74, *p* < .001, Exp. *2*: 𝝌^2^_(1)_ = 29.63, *p* < .001). In line with the third hallmark, high Gaze Prediction trials exhibited a stronger relation between expectation strength and accuracy than low trials (see Fig. S7), and the interaction was significant (*Exp. 1*: mean (SEM) 𝛽 = 0.11 (0.05), *t*_24_ =2.4, *p* = .03; *Exp. 2*: mean (SEM) 𝛽 = 0.18 (0.04), *t*_30_ =4.01, *p* < .001).

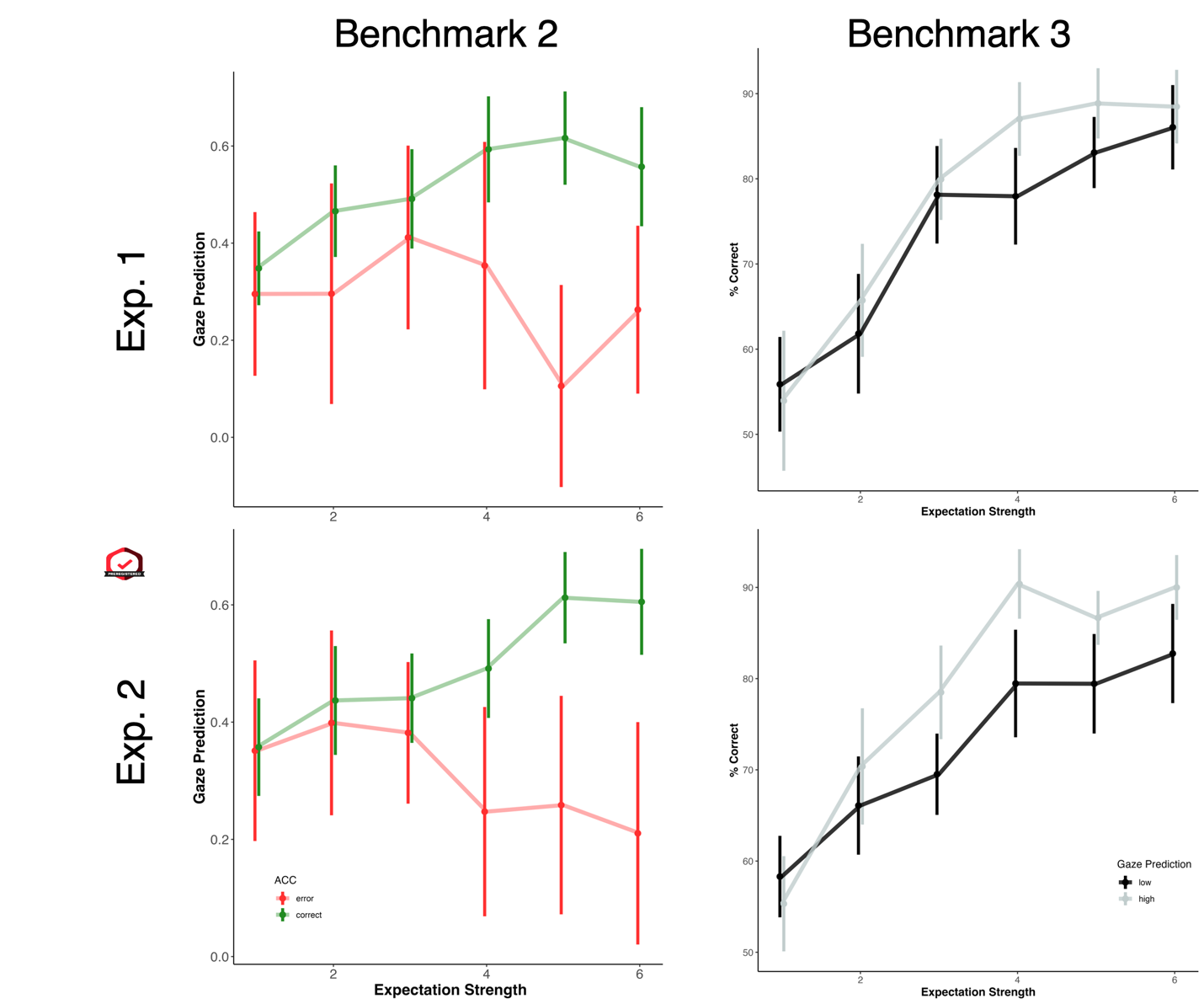

**Figure S7. Gaze Prediction fulfills computational hallmarks of confidence using pre-registered Rescrola-Wagner model.**

##### Simple Effects of Folded-X Pattern

Examining the simple effects of the Folded X interaction, we found both a significant upward slope for correct trials and downward slope for incorrect trials. Noting that there was considerable variability due to some participants having a low number of error trials, we also ran a control analysis including only subjects with at least 30 trials. This corroborated our results and revealed a significant downward slope (Mean β = -.40, SE = 0.16, 𝝌^2^_(1)_ = 6.17, *p* = .01). In addition, we present below the pair-wise comparisons of the different expectation bins (see Table S5 and S6).

| Model | Expectation Bin | *t*_53 =_ | *p* value |
| --- | --- | --- | --- |
| sRW | 1 | 0.32 | 0.75 |
| sRW | 2 | -2.53 | 0.014 |
| sRW | 3 | -3.2 | 0.002 |
| sRW | 4 | -3.07 | 0.003 |
| sRW | 5 | -5.89 | < 0.001 |
| sRW | 6 | -3.89 | < 0.001 |
| RW | 1 | -0.65 | 0.52 |
| RW | 2 | -1.13 | 0.26 |
| RW | 3 | -1.84 | 0.07 |
| RW | 4 | -3.55 | < 0.001 |
| RW | 5 | -6.63 | < 0.001 |
| RW | 6 | -4.29 | < 0.001 |

**Table S5. Pairwise comparison of correct and incorrect trials by expectation strength bin for second hallmark of Gaze Prediction reflecting confidence.** This was performed for both the sRW and pre-registered RW models, combining both experiments.

| model | Expectation Bin | *t*_53 =_ | *p* value |
| --- | --- | --- | --- |
| sRW | 1 | 1.307 | 0.197 |
| sRW | 2 | -2.185 | 0.033 |
| sRW | 3 | -3.752 | < 0.001 |
| sRW | 4 | -2.997 | 0.004 |
| sRW | 5 | -3.555 | 0.001 |
| sRW | 6 | -3.643 | 0.001 |
| RW | 1 | 0.819 | 0.417 |
| RW | 2 | -1.592 | 0.117 |
| RW | 3 | -3.747 | < 0.001 |
| RW | 4 | -4.979 | < 0.001 |
| RW | 5 | -3.367 | 0.001 |
| RW | 6 | -2.354 | 0.022 |

**Table S6. Pairwise comparison of accuracy by expectation strength bin and Gaze Prediction for third benchmark of Gaze Prediction reflecting confidence.** This was performed for both the sRW and pre-registered RW models, combining both experiments.

##### Effect of trial position with block on Folded-X Pattern

To examine how learning influences the Folded X pattern, we analyzed the effect of a trial’s position within each block. Each block was divided into three bins: start/middle/end corresponding to trials at positions 1-12/13-24/15-35. To assess the effect of expectation strength, trials were divided into three equidistant bins based on the decision’s value (see Fig SX). For each participant and each block segment, we fit a separate regression predicting Gaze Prediction from accuracy, expectation strength, and their interaction. The estimate of the interaction was entered into a repeated-measures ANOVA with block segment as the independent variable. This analysis revealed a significant effect of trial position on the interaction (*F*_2,76_ = 8.96, *p* < .001, η^2^ = 0.13; Mean [95% CI] β_Start_ = -.06 [-.18, .06], β_Mid_ = .31 [.14, .48], β_End_ = .40 [.21, .58]). To identify the source of this effect, we examined the slopes for correct and incorrect trials separately. The downward slope for incorrect trials varied significantly across the block and was least pronounced at the start(*F*_2,76_ = 7.16, *p* = .001, η^2^ = 0.11; β_Start_ = .13 [.05, .22], β_Mid_ = -.19 [-.35, .-.23], β_End_ = -.23 [-.41, -.05]), whereas the upward slope for correct trials did not significantly differ by position (*F*_2,96_ = 2.70, *p* = .07). Thus, as learning progressed across the block, the folded X pattern became more pronounced, driven primarily by a steeper downward slope for incorrect trials.

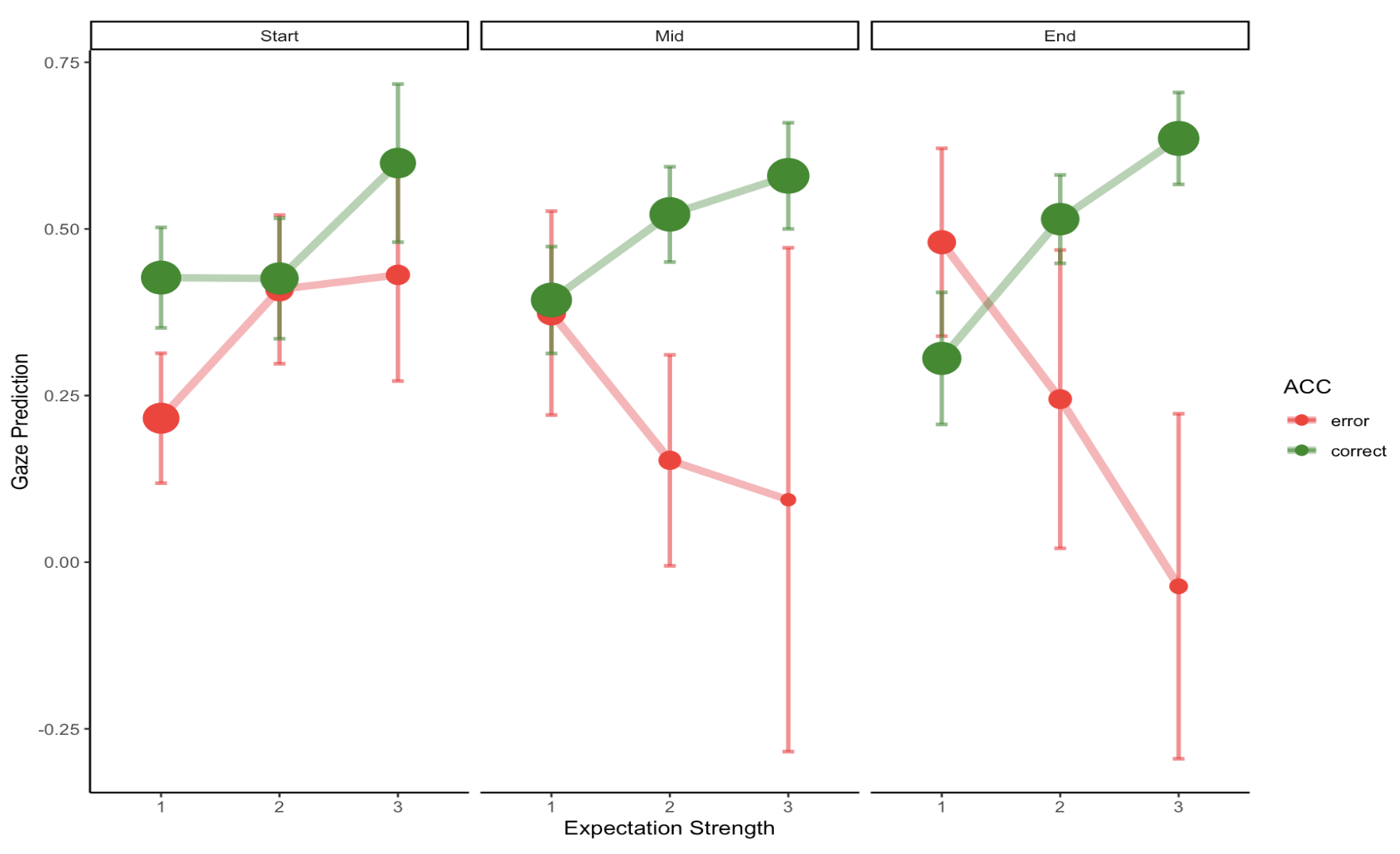

Fig. S8 **Folded X pattern as a function of relative position within a block.** Gaze-based confidence (indexed by Gaze Prediction) is plotted as a function of expectation strength and trial accuracy, separately for trials occurring at the start, middle, and end of each block (positions 1–12, 13–24, and 25–35, respectively). Expectation strength was divided into three equidistant bins, with bin 1 reflecting the lowest expectation strength. Circles represent mean values with 95% confidence intervals; circle size is proportional to the number of trials contributing to each bin across participants.

#### Relation between reported and gaze-based confidence while controlling for additional factors.

To assess the robustness of the finding reported in the main text regarding the association between gaze-based prediction and reported confidence while accounting for additional covariates such as accuracy and the previous trial’s prediction error (see Equation S1). This revealed that the relation between Gaze Prediction and confidence judgments remained significant after controlling for these factors (χ²_(1)_ = 9.5, *p* = .002).

Gaze Prediction ~ confidence rating +accuracy +*p*(Choice *_t_*) +$\delta$*_t_*_-1_ +(1| participant)

(Equation S1)

In addition, we repeated the analysis at the individual-participant level and examined the group-level parameter distribution. Consistent with the main results, reported confidence significantly predicted gaze-based confidence (Mean β = .039, 95% CI [.004, .074], t(31) = 2.30, p = .03, Cohen’s d = 0.41).

Next, because both reported and gaze-based confidence are closely correlated with the model-derived choice probability, we conducted an additional analysis controlling for *p*(Choice *_t_*) (see Equation S2):

Confidence Rating ~ Gaze Prediction + *p*(Choice *_t_*)+ (1| participant)

(Equation S2)

Even after controlling for model-derived choice probability, gaze prediction remained a significant predictor of reported confidence (χ²_(1)_ = 11.62, p < .001). This result was further confirmed when examining the group distribution of the participants’ parameter (Mean $\beta$ = .069, 95% CI = [.021, .12], *t*_31_ = 2.93, *p* = .006, Cohen’s *d* = 0.52).

#### Comparison of reported and gaze-based confidence

##### Correlation of reported and ocular of Metacognition

In line with the results reported in the main text, we found in experiment 3, that both reported confidence ratings and Gaze Prediction exhibited significant metacognitive sensitivity (i.e., Δ confidence > 0), with reported ratings exhibiting significantly higher metacognition. Examining the across subject correlation between the two sources’ metacognition, we did not find a significant correlation, and Bayesian statistics offered anecdotal evidence for the null hypothesis that the two are not correlated (*r_31_* = .09, *p* = .63, BF_10_ = .43; see Fig. S9). The comparison of reported and gaze-based confidence o relates to ongoing debates concerning whether metacognition is a unitary domain-general capacity or comprised of partly dissociable processes (Arbuzova et al., 2023). Although these findings point that the two types of metacognitions are unrelated, they should be interpreted with caution due to the small sample size used (N = 32).

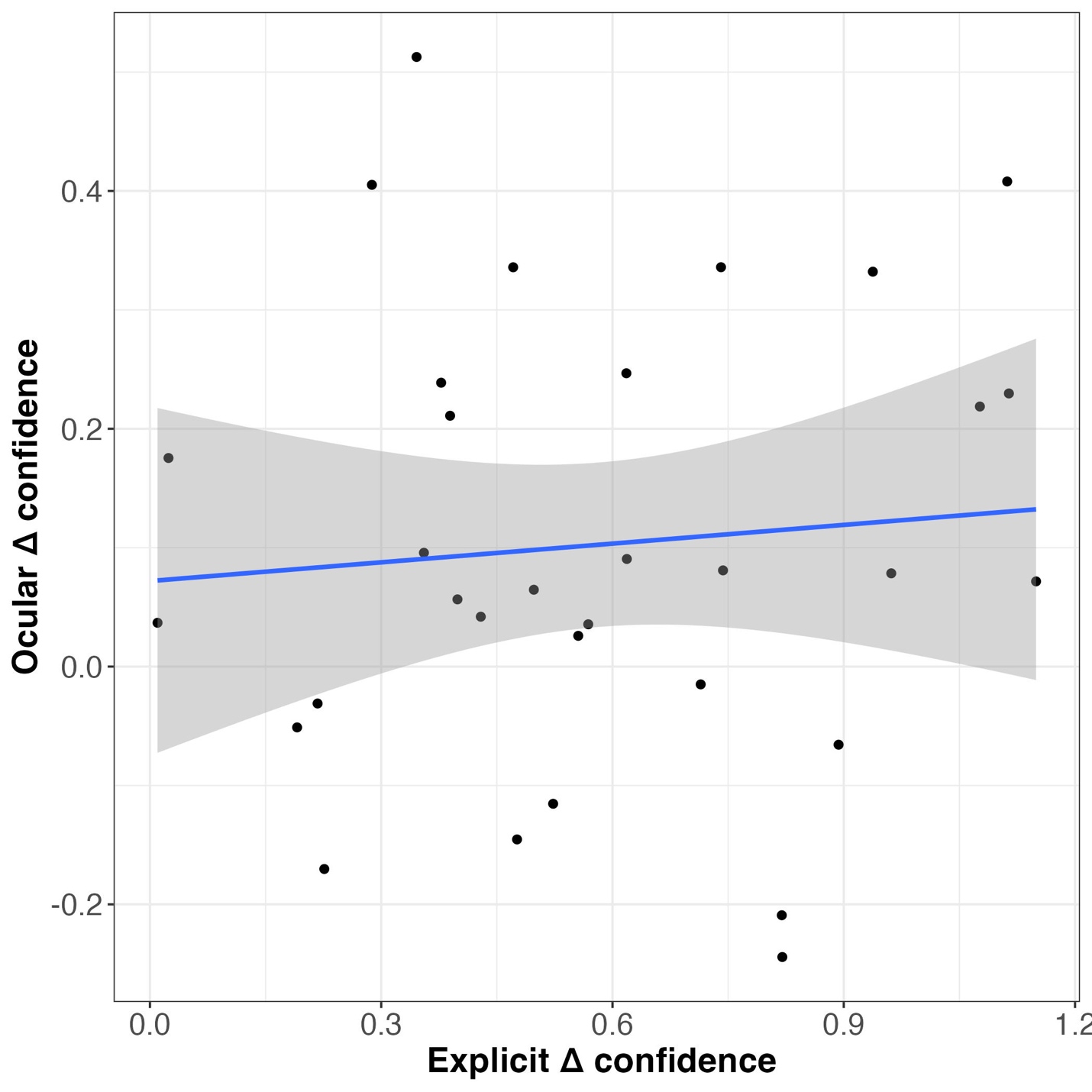

**Figure S9. Correlation of explicit and ocular metacognition across participants in Experiment 3.** In the current sample the correlation was not significant, and Bayesian analysis provided anecdotal evidence for the lack of a correlation. These findings should be interpreted with caution given the small sample size (N = 32).

#### Serial dependence of reported and gaze-based confidence

**Explicit confidence’s serial dependence is independent of additional factors**

To ensure that serial dependence effects are not driven by additional factors such as the previous trial’s accuracy, the current trial’s *p*(Choice *_t_*) probability, we included these factors in our model (see Equation S3). In line with the results reported in the main text, comparing the influence of the previous trial’s confidence rating for reported and gaze-based confidence after controlling for these factors, revealed that reported confidence exhibited significantly greater serial dependence effects (*t*_31_= 6.45, *p* < .001, Cohen’s *d* = 1.14). Thus, the difference found between reported and gaze-based confidence is not driven by additional factors.

Confidence _t_ ~ Confidence_t-1_ + Accuracy_t-1_ + *p*(Choice *_t_*)

(Equation S3)

##### Serial dependence of gaze-based confidence across experiments

Examining serial dependence of gaze-based confidence, we found inconsistent results across the three experiments. In experiment 3, serial dependence for gaze-based confidence was not significantly different than zero (Mean autocorrelation = 0.01, 95% CI = [-.03, .04], *t*_31_= 0.74, *p* = .46). In contrast, gaze-based confidence’s serial dependence across Exp. 1 & 2 was significantly greater than zero (Mean autocorrelation = 0.05, 95% CI = [.02, .07], *t*_56_= 3.40, *p* = .001, Cohen’s *d* =0.45). It remains unclear whether this difference between the experiments stems from statistical power or whether the addition of explicit confidence ratings in experiment 3 reduced ocular confidence’s serial dependence.

##### Serial dependence of reported confidence is extended across trials

To further characterize reported confidence’s serial dependence, we examined whether the autocorrelation is extended beyond adjacent trials (i.e., lag 1). Examining the relation between the current rating and 3 trials back, we found that for lag 2 trial’s confidence was significantly correlated (Mean autocorrelation = 0.04, 95% CI = [.004, .08], *t*_31_= 2.24, *p* = .03, Cohen’s *d* = 0.40), whereas for lag 3 it was not significant (Mean autocorrelation = 0.01, 95% CI = [-.02, .05], *t*_31_= 0.61, *p* = .54).

#### Benefits of convergence and divergence of explicit and ocular learning systems

Exemplifying the benefit of the convergence of the two learning systems we found that accuracy was significantly higher when the explicit and ocular responses converged and were identical) *t*_55_ = 4.75, *p* < 0.001, Cohen’s *d* = 0.63, see Fig. S15).

Examining the benefit of the divergence of the two responses types we found that exploratory trials (i.e., *p*(Choice *_t_*) < .5) exhibit a significantly higher divergence rate of explicit and ocular responses(*t*_53_ = 4.13, *p* < 0.001, Cohen’s *d* = 0.56, see Figure S10).

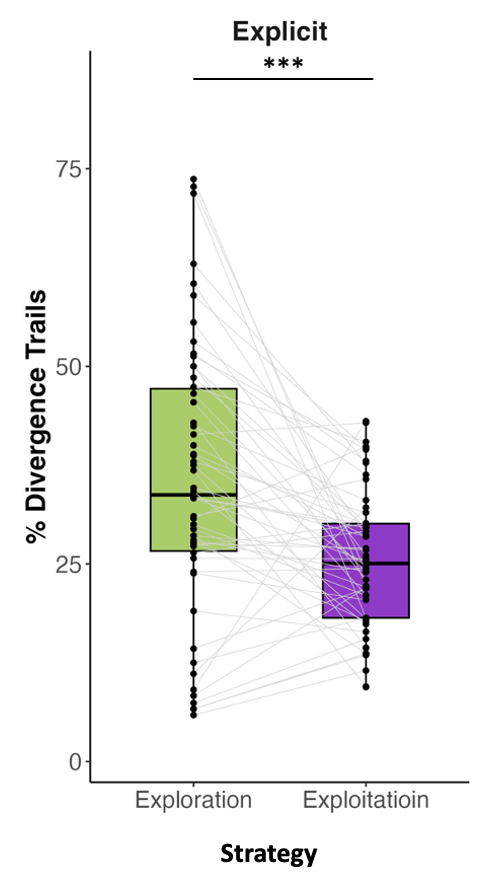

**Figure S10. Increased divergence of explicit and ocular responses on exploratory trials.** Responses diverged significantly more on exploratory trials (i.e., p(Choice *_t_*) < .5) as opposed to exploitation trials (i.e., p(Choice *_t_*) > .5). p(Choice *_t_*).

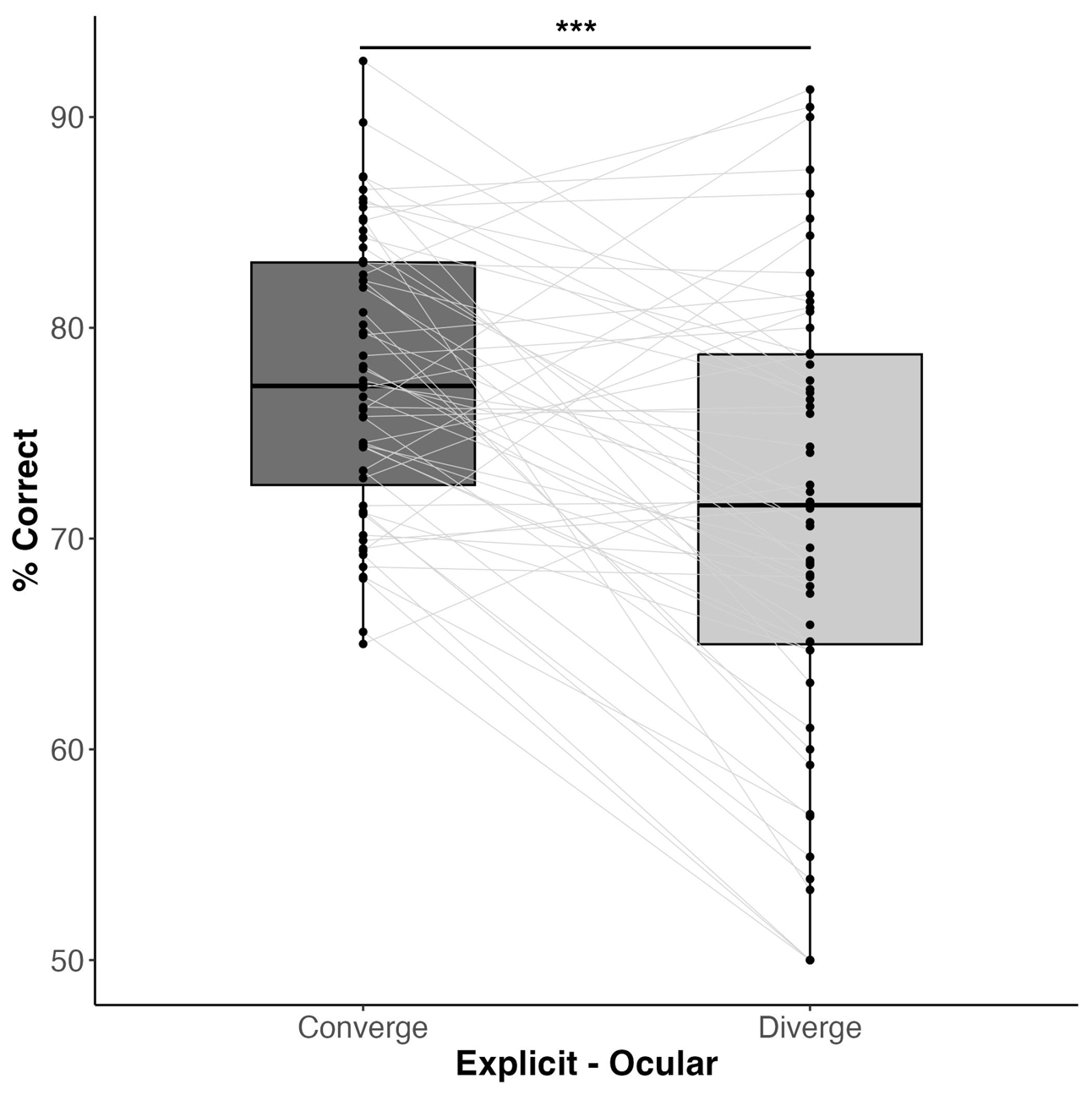

**Figure S11. Explicit responses rule accuracy on converging trials is significantly higher than on diverging trials.** Converging trials are trials in which the explicit response and binarized ocular ‘response’ are identical (e.g., both are “right”).

#### Additional Pre-Registered Analyses of Experiment 3

In the current section we report the results of the additional analyses that were pre-registered for experiment 3 (<https://osf.io/ngsx6>). To recap, in experiment 3 we examined the relation between explicit ratings of confidence and ocular-derived confidence-like behavior. Participants performed the same task as in previous experiments, with the addition that following their prediction of the target’s location they rated their confidence on a five-point vertical slide. In the current section we report the additional analyses (hypothesis 3&4) that were not central to the results in main text.

##### Hypothesis 3: Explicit confidence ratings track learning processes

This was composed of two sub-hypotheses. First, we tested this using model free tests (i.e., pre-registered H3a). In line with our first test, we found that confidence ratings significantly decrease following an error (Mean difference of confidence for correct vs. incorrect trials [95% CI] = 0.79 [0.61, 0.98], *t*_31_ = 8.73, *p* < 0.001, Cohen’s *d* = 1.54, see Fig. S12A), exhibiting a core feature of learning (Miller et al., 1995; Rescorla, 1972). Furthermore, and in line with our second test, normalized confidence ratings exhibited a learning curve (Fig. S12B), further supporting confidence’s convergence with learning processes. Finally, in line with our additional test (i.e., pre-registered H3b), confidence exhibited metacognitive sensitivity and delta confidence was significantly greater than zero (see Fig. 4b and main text for statistical result).

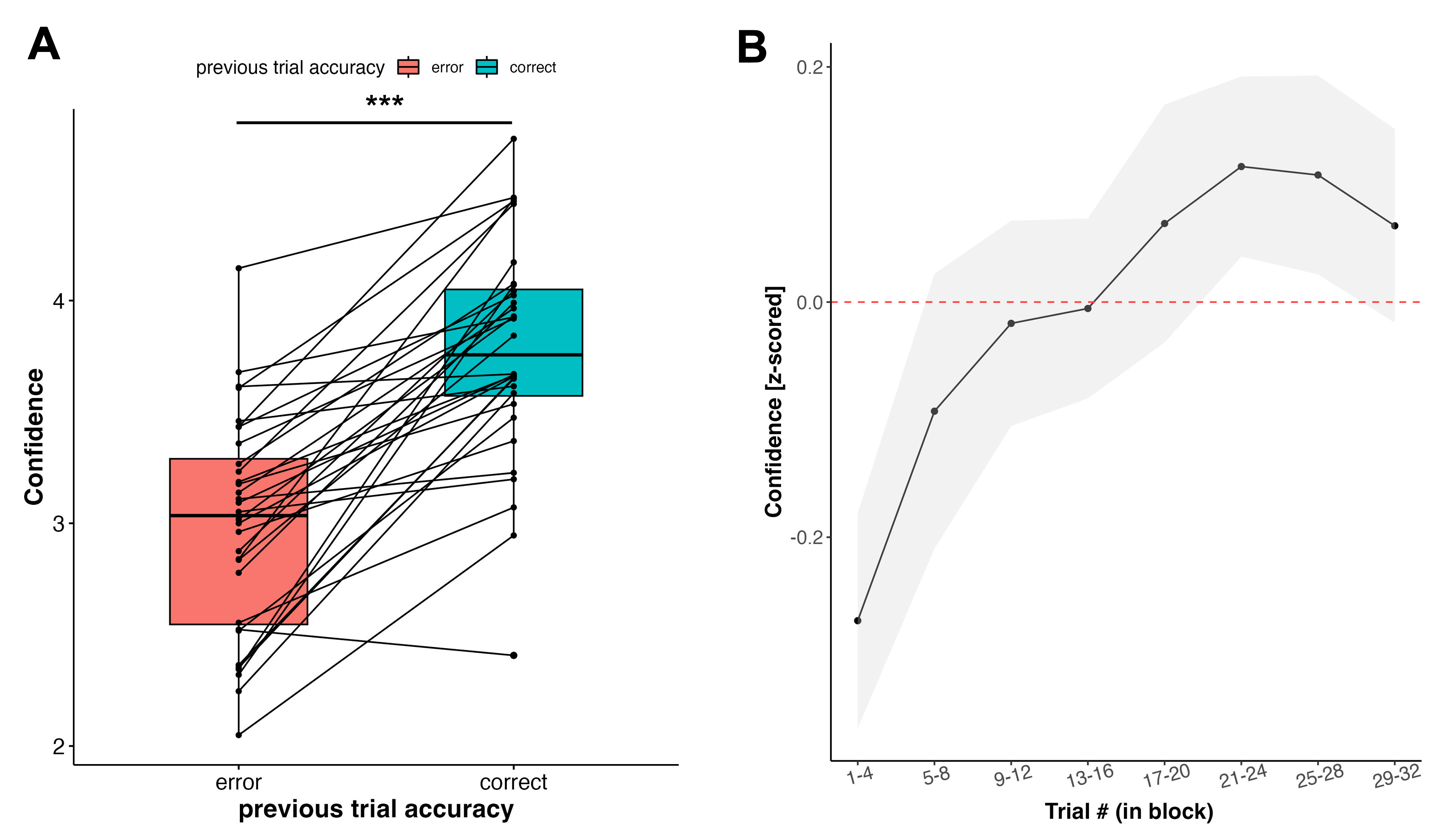

**Figure S12. Confidence exhibits model-free characteristics of learning. (A)** Confidence was significantly decreased following erroneous trials in contrast to correct trials. **(B)**. Normalized confidence ratings across blocks and participants exhibited a learning curve. At the start of the block, confidence gradually increases until it reaches a plateau towards the final third of the block. Grey area represents 95% CI across participants. Individual participants learning curve was calculated across blocks.

Next, utilizing model-based metrics of learning (i.e., pre-registered Hypothesis 3b), we examined the trial-by-trial correlation between confidence ratings and computationally derived quantities: the previous trial’s prediction error (δ_t-1_), and the probability of the choice made (*p*(Choice *_t_*)). In line with our pre-registration both were significantly correlated to confidence ratings (δ_t-1_: Mean Mean group $\rho$ [95% CI] = -.27 [-.19, -.34], *t*_31_ = 7.6, *p* < .001, Cohen’s *d* = 1.35, 22/32 participants significantly correlated at individual level; *p*(Choice *_t_*): Mean Mean group $\rho$ [95% CI] = .38 [.33, .44], *t*_31_ = 14.5, *p* < .001, Cohen’s *d* = 2.56, 30/32 participants significantly correlated at individual level; see Fig. S13). Thus, in line with previous work (Hertz et al., 2018; Meyniel et al., 2015), confidence ratings closely track the computationally derived probability of the choice made.

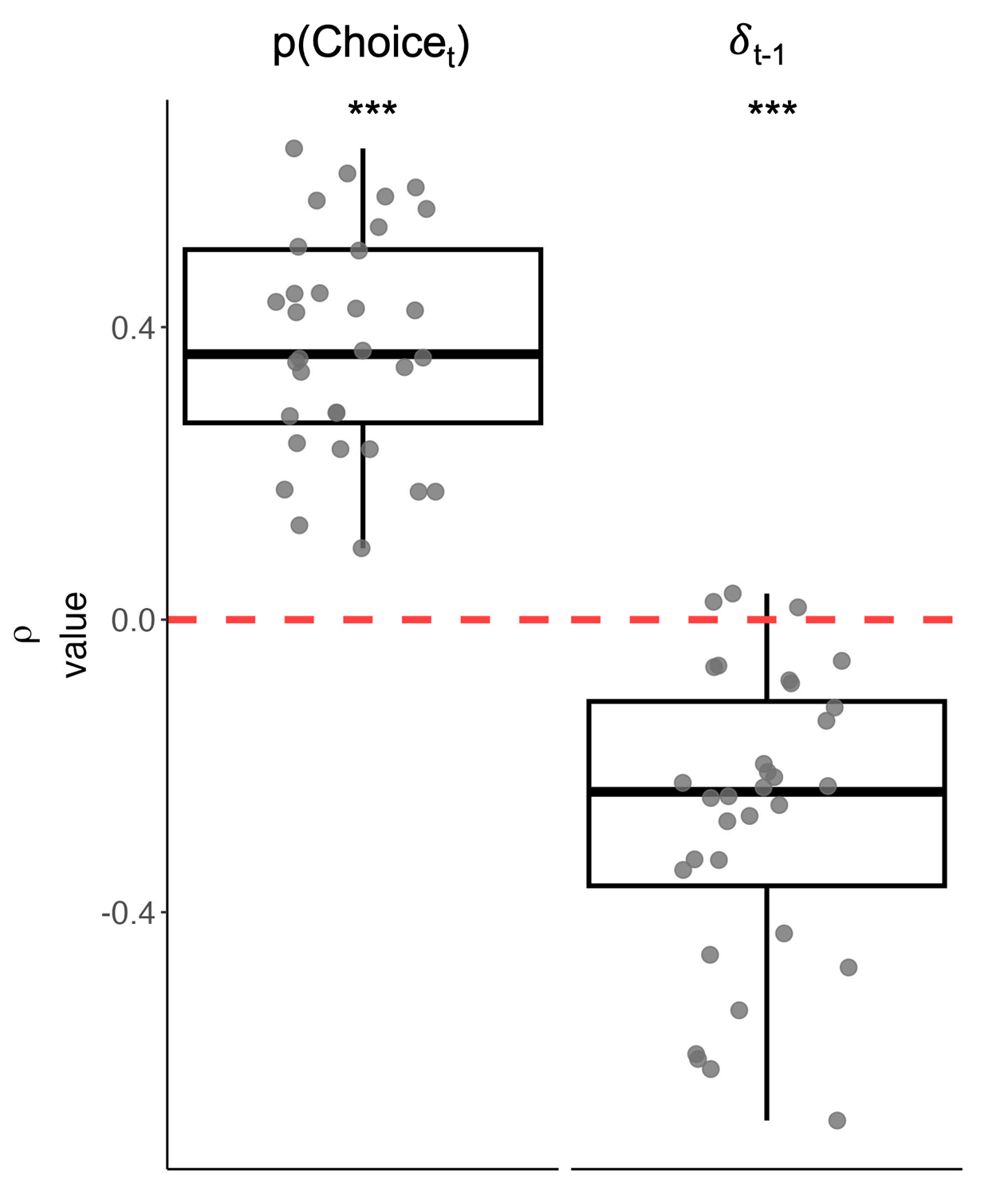

**Figure S13. Explicit confidence ratings closely track latent learning variables.** Trial-by-trial confidence ratings were correlated with the latent learning variables: the underlying probability of the choice made (i.e., *p*(Choice *_t_*)) and the previous trial’s prediction error (i.e. δ_t-1_) were calculated per participant. The group distribution of the Pearson coefficients is presented above, and as pre-registered is highly significant for both. Thus, confidence ratings track the underlying learning processes.

##### Hypothesis 4: Replication of gaze’s close relation to explicit predictions.

In line with our robust findings linking ocular expectations to explicit predictions in the first two experiments, we sought to replicate these findings in Exp. 3. Importantly, Exp. 3 expands our previous findings by demonstrating the link between gaze and explicit responses in an experimental paradigm in which a confidence question separates their response and the period in which gaze is measured.

In line with our pre-registration, we found a robust link between gaze and explicit predictions. Specifically, in line with hypothesis 4A, Gaze Direction was significantly more in the direction of the explicit prediction (Mean gaze difference = 0.90, 95% CI= [0.75, 1.04], *t*_31_ = 12.79, *p* < .001, Cohen’s *d* = 2.26; see Fig. S14A). Furthermore 30 out of 32 participants exhibited a significant effect at the individual level (see Fig. S14B). Thus, despite the interruption of the confidence question, gaze was still closely aligned with the explicit prediction’s direction in a magnitude like that of the previous experiments.

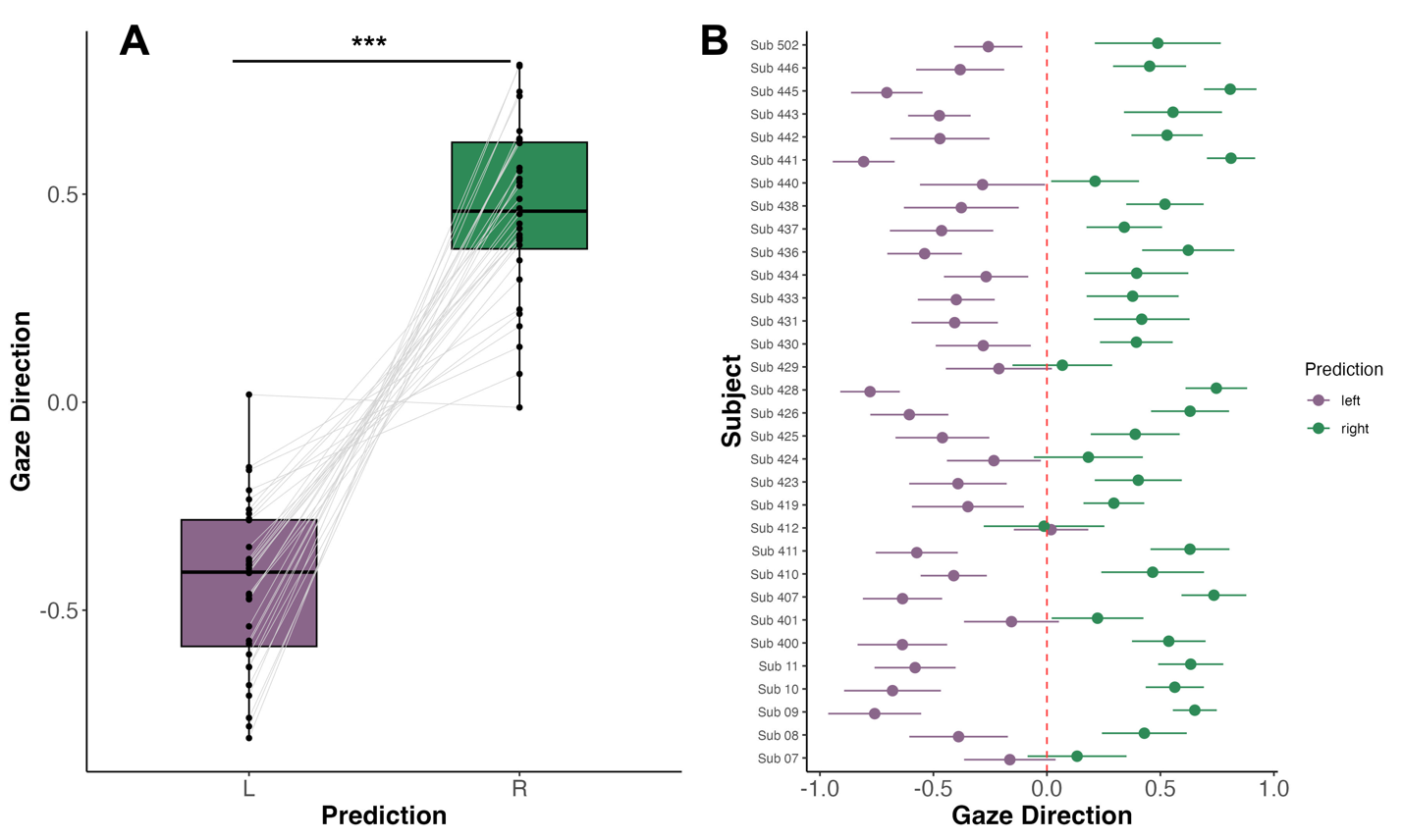

**Figure S14. Gaze direction closely follows explicit prediction despite addition of confidence question in Exp. 3. (A)** Gaze direction was significantly aligned with the explicit prediction’s direction. **(B)** At the individual subject level, all participants except for two exhibited a significant effect.

Turning to examine Gaze Prediction’s relation to the underlying probability of the choice made (i.e., *p*(Choice *_t_*)), we examined the trial-by-trial correlation between the two. In line with previous experiments the group distribution of correlation coefficients was significantly greater than zero (Mean $\rho$= .07, 95% CI = [.037, .10], *t*_31_ = 4.26, *p* < .001, Cohen’s *d* =0.75; see Fig. S15). It should be noted that although significant the magnitude of correlation is smaller than the previous experiments mean correlation coefficient (Exp. 1: Mean $\rho$= .13 [.08, .19], Exp. 2: Mean $\rho$= .11 [.05, .16]). Thus, the introduction of the confidence rating reduces this link between Gaze Prediction and *p*(Choice *_t_*).

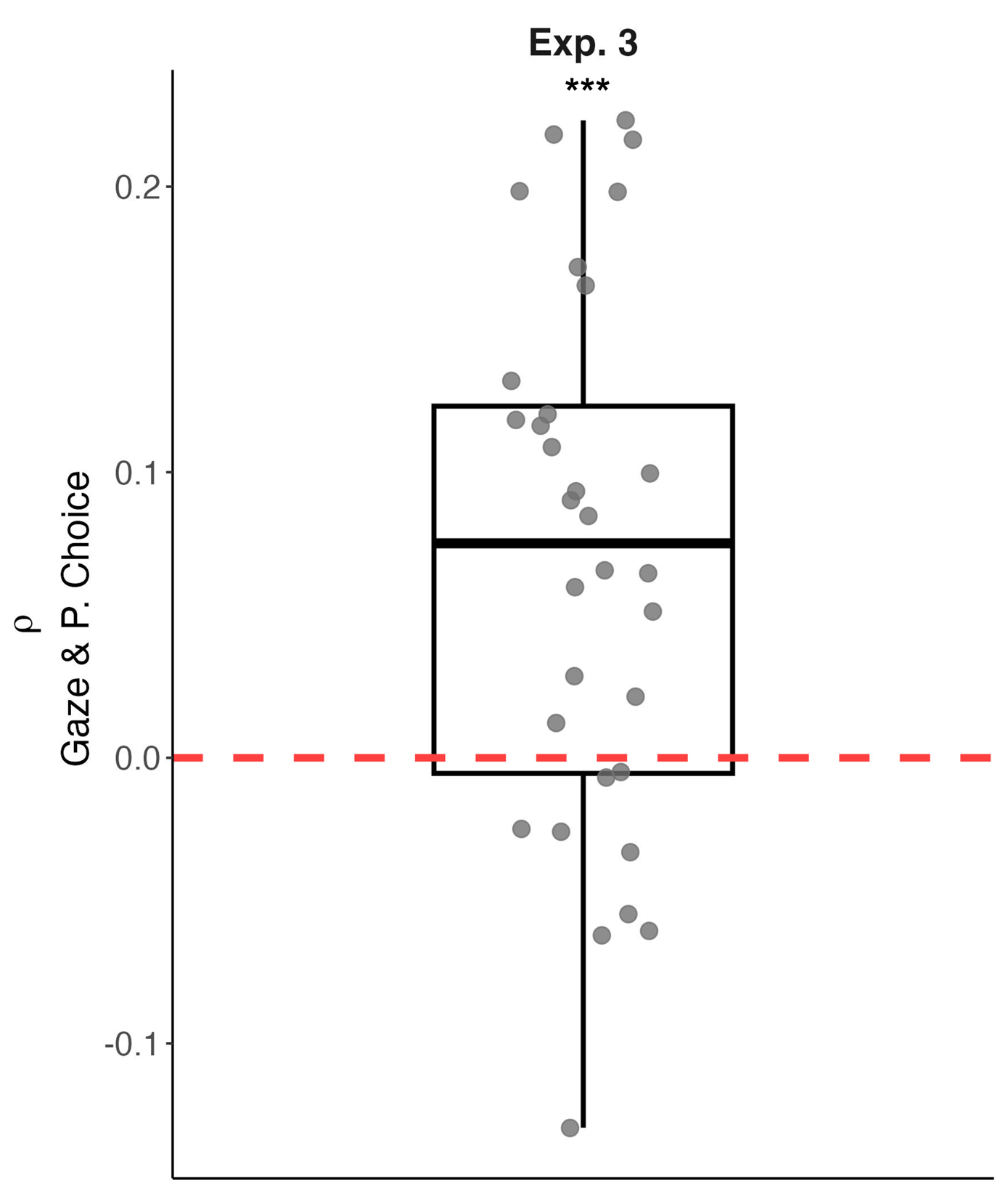

**Figure S15. Correlation of Gaze Prediction and *p*(Choice*_t_*) in Exp. 3.**

##### Gaze Prediction partially fulfills computational benchmarks of confidence in Exp. 3

Turing to the pre-registered hypothesis 4B we examined whether Gaze Prediction exhibits the three computational benchmarks of confidence (Sanders et al., 2016), and using the same analysis plan described in the main text and in the pre-registration. Regarding the first benchmark, we found a positive monotonic relation between accuracy and Gaze Prediction. The group’s correlation coefficient distribution was showed a non-significant trend towards being greater than zero (Mean $\rho$ = .18, 95% CI = [-.01, . 63], *t*_31_ = 1.97, *p* = .057, Cohen’s *d* =0.35; see Fig S16A). Importantly, examining this relation via a more rigorous linear mixed model (see Equation S4) that accounts for individual difference we found a significant effect of Gaze Prediction on *p*(Choice *_t_*) (𝝌^2^_(1)_ = 5.70, *p* < .001). Examining the second benchmark of a folded X pattern, we did not find a significant interaction of *v* and the trial’s accuracy (𝝌^2^_(1)_ = 1.91, *p* = .17; see Fig. S16B). Examining the third benchmark, high Gaze Prediction trials exhibited a stronger relation between *v* and accuracy than low gaze trials (see Fig. S16C), and the interaction was significant (mean (SEM) 𝛽 = 0.08 (0.04), *t*_31_ =2.02, *p* = .04;). To summarize, Gaze prediction fulfilled the first and third benchmark, whereas the second was not replicated. Similar to the reduced correlation between gaze and *p*(Choice *_t_*), gaze’s does not fully fulfill the benchmarks of confidence. It may be that the introduction of the confidence rating introduces additional cognitive demands that in turn reduce gaze’s relation to confidence. An additional possibility is that the explicit rating of confidence alters the confidence-like calculations that gaze performs resulting in a partial fulfillment of the confidence benchmarks.

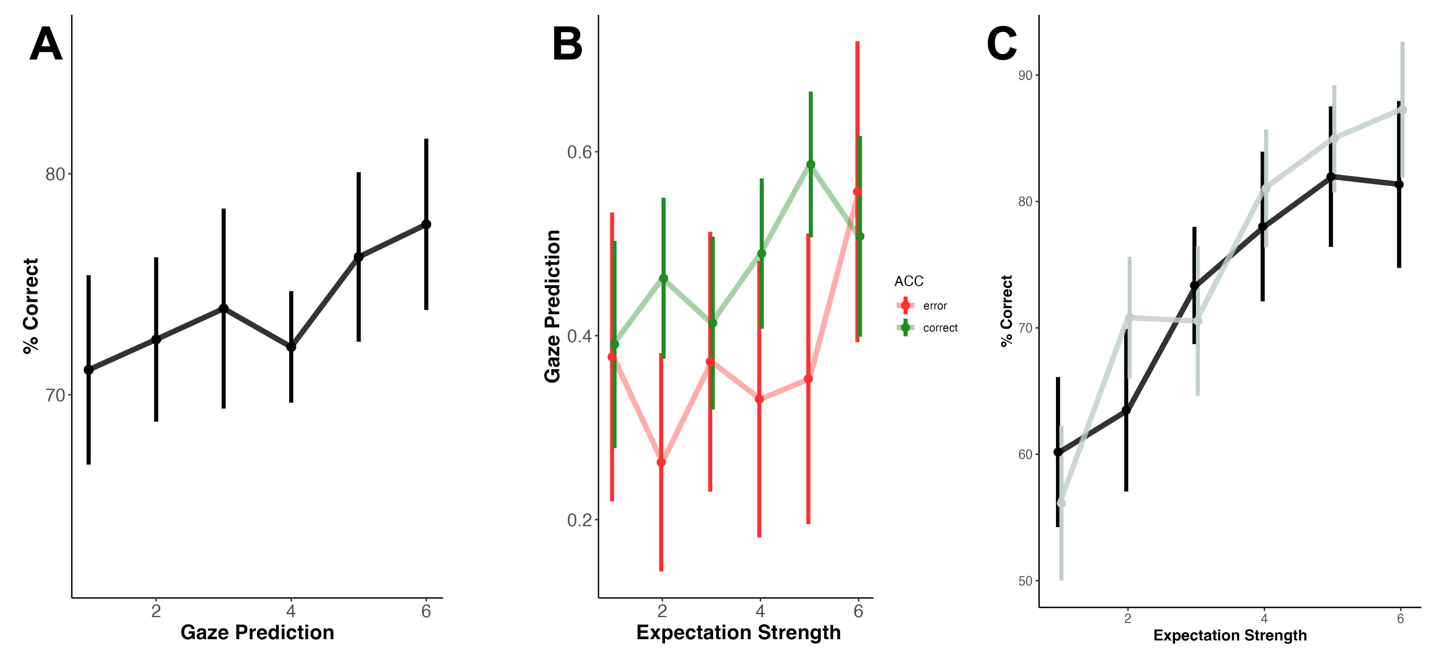

**Figure S16. Gaze partially fulfills computational benchmarks of confidence in Exp. 3.** In Exp. 3, following the explicit prediction participants explicitly rated their confidence on a vertical scale, and after which they entered the virtual environment. Importantly, gaze partially fulfilled the computational benchmarks of confidence, further expanding our findings in the first two experiments. **(A)** There was a significant monotonic rise in accuracy as Gaze Prediction increased. **(B)** ‘Folded X pattern’, in contrast to the previous experiments, Gaze Prediction did not fulfill this benchmark. **(C)** Trials with high Gaze Prediction showed a significantly stronger relation between expectation strength and accuracy.

Acc ~ Gaze Prediction+(1|participant)

(Equation S4)

### Methods

#### Experimental paradigm

##### Hardware and Setup

The experiment was conducted on a high performance desktop (CPU: 11th generation Intel Core i9 processor, RAM: 64 GB, GPU: NVIDIA GeForce RTX 3080) running in-house software (developed using Unity 2018.3.2). An HTC Vive Pro Eye head mounted display (HMD), Vive controller, and two base stations were used to track participants' hand motions and generate the virtual environment projected through the HMD (refresh rate ~ 90 Hz). The trackpad of the Vive controller served as an input device to record participant’s explicit prediction of the upcoming target’s location (Fig. 1A). Eye tracking was performed via the built-in Tobii trackers at a sampling rate of ~90 Hz, and extracted using the HTC SRanipal SDK.

The participants were seated at a table, while wearing the HMD and holding the controller in their right hand. They placed the controller on a textured elevated black 'x' sign in the physical environment that mirrored the ‘x’ in the virtual environment. The ‘x’ served as the trial’s beginning and end point. The ‘x’ was slightly elevated and textured, enabling participants to find it via touch while wearing the VIVE HMD. They controlled a virtual hand that was tracked via the controller. The virtual hand looked and moved similarly to their real hand in space and time.

After completing the probabilistic learning task, participants also performed a sensorimotor conflict-detection task to assess their sense of agency (Applebaum et al., 2025) and completed several self-report questionnaires pertaining to psychotic-like and anomalous self-experiences (Cicero et al., 2017; Foa et al., 2002; Loewy et al., 2011). These data were collected for exploratory purposes and are not included in the present article.

##### Trial Sequence

The number of trials within each block and whether all participants were presented with identical trial sequences varied slightly between experiments. Specifically, in Exp. 1 each block consisted of 32 trials, and each participant was presented with a unique pseudo-random trial sequence. In Exp. 2, trial sequence was fixed across participants and block length was between 29-35 trials, with a rule switch occurring at trials: 33, 64, 99, and 132. In this experiment we used a fixed sequence following previous computational works (Harrison et al., 2021; Iglesias et al., 2013)h to reduce noise arising from variance in stimuli order and to enhance the comparability of model parameters across participants. In Exp. 3, block length was like Exp. 2, yet each participant was presented with a unique pseudo-random trial sequence.

#### Measures

##### Gaze Prediction transformation

To obtain a measure that reflects the congruence of gaze direction and the explicit prediction, we normalized the gaze (that is normalized per subject across all trials in the 300 ms interval) by the explicit prediction using the following algorithm:

$$Gaze Prediction=\left\{ \begin{aligned} G\mathrm{aze}\mathrm{Normalized} if Prediction ="Right" \\ G\mathrm{aze}Normalized*-1 if Prediction ="Left" \end{aligned} \right.$$

(Equation S5)

Thus, positive values reflect Gaze in the direction of the prediction (regardless of its actual direction), and negative values reflect gaze in the opposite direction of the explicit prediction.

##### Analysis plan for examining whether Gaze Prediction fulfills computational benchmarks of confidence.

The analysis plan and its justification are described in detail in the pre-registration (https://osf.io/p4hyq). As stated in the pre-registration: “To adapt the theoretical framework of Sanders et al., (2016) that was originally applied to tasks from the signal detection framework (e.g., tone detection) we made three changes in our analyses. First, instead of utilizing accuracy (did the participants prediction match the target’s actual location), we examined rule accuracy (i.e., did the participants' response abide by the underlying rule). Second, as a proxy for evidence we utilized the trial’s expectation strength (𝓋) derived from the Rescorla Wagner model. Following the contingency coding in the modeling (Iglesias et al., 2013), this was formulized as the expectation’s distance from 0.5, and was binned into six percentiles for each participant. Finally, instead of examining confidence rating we examine the ocular measure of Gaze Prediction.”

#### Model fitting of learning

To analyze learning dynamics throughout the experiment, we employed a model-based approach based on The model space consisted of: (1) the well-established Rescorla-Wagner (RW) model (Rescorla, 1972); (2) a Rescorla-Wagner model with an additional stickiness parameter (sRW) (Culbreth et al., 2016; Daw et al., 2011); and (3) a noisy Win-Stay Lose-Switch (WSLS) model (Wilson & Collins, 2019).

The RW model consists of a perceptual and decision model. In the perceptual model each option’s value (*v*) is updated by the prediction error (δ)—the discrepancy between the previous trial’s predicted and actual outcomes (*r*)—modulated by the learning rate (𝛼), a free parameter that controls the extent of updating by prediction errors (Eq. S6A). Lower 𝛼 values suggest slower learning over more trials. The value (*v*) from the perceptual model informs the decision model, which employs a SoftMax function to calculate the choice probability (i.e., *p*(Choice *_t_*)). The function’s steepness and randomness are dictated by the inverse temperature (β), another free parameter, reflecting decision consistency (Eq. S6B).

(Eq. S6A)	$v_{t+1}=v_{t}+ \alpha*\delta_{t}$

Where	$\delta_{t}= r_{t}-v_{t}$

(Eq. S6B)	${\boldsymbol{P}.}_{Choice}=\frac{e^{\beta*\boldsymbol{v}}}{e^{\beta*\boldsymbol{v}}+e^{\beta*(1-\boldsymbol{v})}}$

The "sticky" sRW model extends the traditional RW model by adding a perseveration parameter (⍴) to quantify the tendency to use the rule used in the previous decision (Culbreth et al., 2016; Daw et al., 2011). This parameter is included in the decision model (Eq. S7), multiplied by *rep (a)*, an indicator function that equals 1 for decisions that apply the rule used in the previous decision and 0 otherwise. Importantly, decisions are coded in a cue-contingent manner (cue A and a “right” decision are coded identically to cue B and a “left” response), which implies that perseveration reflects that a decision adheres to the same rule across trials, rather than mere motor repetition. Notably, the RW model is nested within the sRW model when the parameter ⍴ is set to zero.

Eq. S7	${P.}_{Choice}=\frac{e^{\beta*\boldsymbol{v+}⍴*rep(a)}}{e^{\beta*\boldsymbol{v+}⍴*rep(a)}+e^{\beta*\left( 1-\boldsymbol{v} \right)+ ⍴*rep(a)}}$

Unlike the RW model, which uses a free parameter to determine the integration window, in the WSLS model (Eq. S8) decisions are based solely on the previous trial's outcome, and a free parameter (𝜺) incorporates probabilistic noise in the decision-making.

Eq. S8	${P.}_{Choice}\left\{ \begin{aligned} 1-\varepsilon if C_{t-1} correct \\ \varepsilon if C_{t-1} incorrect; \end{aligned} \right.$

For all models, stimuli and responses were encoded within a unified contingency space to minimize free parameter estimates, in line with our pre-registration and previous work (Harrison et al., 2021; Iglesias et al., 2013). Model parameters were estimated via the Bayesian adaptive direct search (BADS) toolbox (Acerbi & Ji, 2017), optimizing each model's log likelihood. Parameter recovery across models was reliable (see Fig. S19). Model comparison was performed via the Bayesian Information Criteria (BIC; (Schwarz, 1978)). All models showed good model recovery (see Fig. S20).

##### Use of delta confidence as measure of metacognitive sensitivity

The analysis plan and its justification are described in detail in the pre-registration (https://osf.io/ngsx6). To obtain an ocular “confidence” rating that is comparable measure to participant’s confidence rating that is performed on a 5-point scale, Gaze Prediction was binned into five equidistant bins ranging from the minimum to maximum value per participant. To compare the measures’ metacognition, we choose delta confidence because it is a measure with few assumptions and high interpretability. It should be noted that first-order performance is identical for the explicit and ocular confidence given that each rating is obtained for each trial. Hence there is no need for measures such as meta-d’ that attempt to control for differences in first order performance and include additional assumptions.

#### Computational modeling

##### Parameter estimation procedure

Computational modeling was done following the model logic used in previous works (Harrison et al., 2021; Iglesias et al., 2013) and the Hierarchical Gaussian Filter toolbox (Frässle et al., 2021). To reduce the number of free parameters, stimuli and responses were coded in a single contingency space resulting in the modeling of a single learning trajectory (Iglesias et al., 2013). Importantly, the values of the contingency coding (0/1) are arbitrary so accordingly we utilize the distance from v = 0.5 as the strength of the value assigned and the unsigned prediction error to assess prediction errors.

Model fitting was done using the Bayesian adaptive direct search (BADS; (Acerbi & Ji, 2017)) toolbox to optimize the log likelihood of each model and in-house scripts. Parameters’ plausible search space was defined based on the distribution of values typically found in the literature and a deterministic search was used. Each subject’s data was fit separately ten times using different random initial starting points drawn from the plausible bounding box. The fitting iteration that yielded the best likelihood and converged, its parameters were chosen and inserted to group level analyses. For external validation we compared these model parameters to those using the TAPAS HGF toolbox,and the two obtained similar parameter estimates.

##### Parameter recovery

Parameter recovery was performed for all models across 1,000 iterations. All parameters showed good to reasonable recovery (see Fig. S17 & Table S8) that were highly significant. As can be seen in the figure, extreme values showed lower recovery rates, and occasionally recovered estimates were mistakenly estimated as extreme values. In line with these limitations that were especially prominent for β, in the analyses presented we treat with caution extreme values (i.e., β > 15) that were estimated.

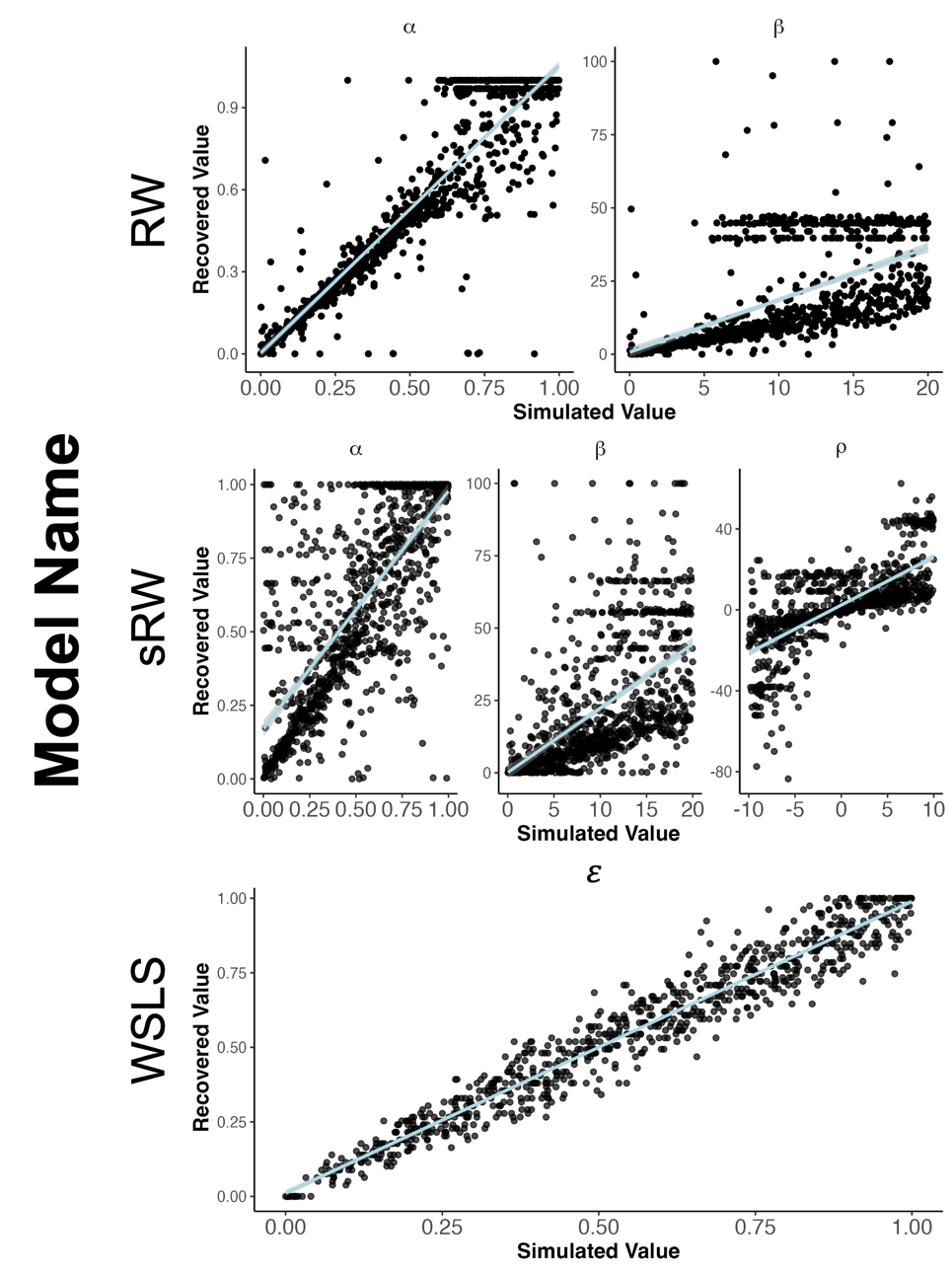

**Figure S17.Parameter recovery across models.** Parameter recovery was done by simulating 1,000 iterations of values uniformly distributed across the plausible interval of each parameter. Across all parameters there was reasonable to high recovery rates. As seen in the recovery of β, high values were occasionally mistakenly obtained. Accordingly in the parameters obtained from the actual data we treat these values with caution.

| Model Name | Parameter | Mean correlation value [95% CI] |
| --- | --- | --- |
| RW | alpha | 0.92 [0.91, 0.93] |
| RW | beta | 0.59 [0.54, 0.63] |
| sRW | alpha | 0.73 [0.70, 0.76] |
| sRW | beta | 0.57 [0.53, 0.61] |
| sRW | rho | 0.71 [0.67, 0.74] |
| WSLS | epsilon | 0.97 [0.97, 0.98] |

**Table S8. Correlation between simulated and recovered parameters.** All parameters showed significant correlations.

##### Model Recovery

Model recovery was performed across 1,000 iterations and showed good recovery (see Figure (S18).

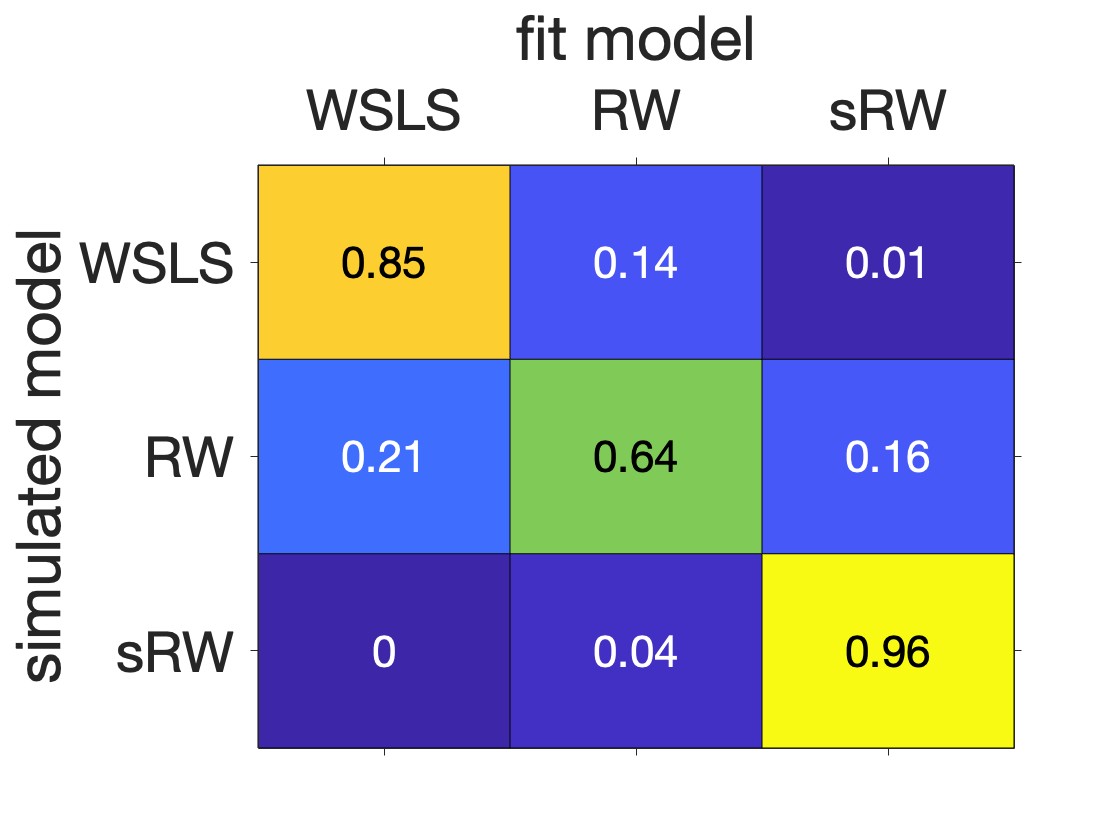

**Figure S18. Model recovery for models.** The model that simulated the data (vertical axis) was reliably recovered (horizontal axis). Values in box signify the percentage that the model was fit per each simulated row (i.e., each row’s sum is one). As can be seen the diagonal shows the highest values signifying that the simulated model was in large recovered.
